# Immunogenicity and protective efficacy of one- and two-dose regimens of the Ad26.COV2.S COVID-19 vaccine candidate in adult and aged rhesus macaques

**DOI:** 10.1101/2020.11.17.368258

**Authors:** Laura Solforosi, Harmjan Kuipers, Sietske K. Rosendahl Huber, Joan E.M. van der Lubbe, Liesbeth Dekking, Dominika N. Czapska-Casey, Ana Izquierdo Gil, Miranda R.M. Baert, Joke Drijver, Joost Vaneman, Ella van Huizen, Ying Choi, Jessica Vreugdenhil, Sanne Kroos, Adriaan H. de Wilde, Eleni Kourkouta, Jerome Custers, Tim J. Dalebout, Sebenzile K. Myeni, Marjolein Kikkert, Eric J. Snijder, Dan H. Barouch, Kinga P. Böszörményi, Marieke A. Stammes, Ivanela Kondova, Ernst J. Verschoor, Babs E. Verstrepen, Gerrit Koopman, Petra Mooij, Willy M.J.M. Bogers, Marjolein van Heerden, Leacky Muchene, Jeroen T.B.M. Tolboom, Ramon Roozendaal, Hanneke Schuitemaker, Frank Wegmann, Roland C. Zahn

**Affiliations:** Janssen Vaccines and Prevention B.V., Leiden, The Netherlands; Molecular Virology Laboratory, Department of Medical Microbiology, Leiden University Medical Center, Leiden, The Netherlands; Center for Virology and Vaccine Research, Beth Israel Deaconess Medical Center, Harvard Medical School, Boston, MA 02215, USA; Biomedical Primate Research Centre, Rijswijk, The Netherlands; Non-Clinical Safety Toxicology/Pathology, Janssen Research and Development, Beerse, Belgium

## Abstract

Safe and effective coronavirus disease (COVID)-19 vaccines are urgently needed to control the ongoing pandemic. While single-dose vaccine regimens would provide multiple advantages, two doses may improve the magnitude and durability of immunity and protective efficacy. We assessed one- and two-dose regimens of the Ad26.COV2.S vaccine candidate in adult and aged non-human primates (NHP). A two-dose Ad26.COV2.S regimen induced higher peak binding and neutralizing antibody responses compared to a single dose. In one-dose regimens neutralizing antibody responses were stable for at least 14 weeks, providing an early indication of durability. Ad26.COV2.S induced humoral immunity and Th1 skewed cellular responses in aged NHP that were comparable to adult animals. Importantly, aged Ad26.COV2.S-vaccinated animals challenged 3 months post -dose 1 with a SARS-CoV-2 spike G614 variant showed near complete lower and substantial upper respiratory tract protection for both regimens. These are the first NHP data showing COVID-19 vaccine protection against the SARS-CoV-2 spike G614 variant and support ongoing clinical Ad26.COV2.S development.

**Summary:** COVID-19 vaccines are urgently needed and while single-dose vaccines are preferred, two-dose regimens may improve efficacy. We show improved Ad26.COV2.S immunogenicity in non-human primates after a second vaccine dose, while both regimens protected aged animals against SARS-CoV-2 disease.

## Introduction

Development of safe and effective vaccines to control the ongoing COVID-19 pandemic (Cucinotta and Vanelli, 2020; WHO, 2020) caused by severe acute respiratory syndrome coronavirus 2 (SARS-CoV-2) (Wu et al., 2020; Zhu et al., 2020) is a global priority. Ideally, especially in the context of a pandemic, a vaccine provides both an early onset of protection and durable protection. The durability of vaccine-elicited protection depends on the capacity of the vaccine platform, specific antigen (design) and vaccination regimen to efficiently stimulate the immune system (Cohen, 2019; Pulendran & Ahmed, 2011) and on several characteristics linked to the recipient of the vaccine (Zimmermann and Curtis, 2019). Age for instance plays an important role, as in the elderly the immune response to vaccines is usually reduced in magnitude and duration, potentially resulting in reduced vaccine efficacy (Wagner et al., 2018; Crooke et al., 2019; Gustafson et al., 2020; Weinberger, 2018). Although persons of all ages are at risk of contracting COVID-19, the risk of developing severe or critical illness increases markedly with age (Mallapaty, 2020; CDC, 2020), warranting the testing of COVID-19 vaccine candidates in different age cohorts.

The Ad26.COV2.S vaccine candidate is a non-replicating adenovirus 26 (Ad26)-based vector encoding the stabilized full length SARS-CoV-2 spike protein based on the Wuhan Hu1 SARS-CoV-2 isolate, and containing an aspartic acid (D) residue in amino acidic position 614 (D614) (Bos et al., 2020). In pre-clinical efficacy studies, a single dose of Ad26.COV2.S, provided robust protection against SARS-CoV-2 challenge (USA-WA1/2020 viral strain, D614) in both upper and lower airways in rhesus macaques (Mercado et al., 2020) and protected Syrian golden hamsters from severe clinical disease (Tostanoski et al., 2020). Protective efficacy against SARS-CoV-2 in NHP in this and other studies, strongly correlated with the presence of virus binding and neutralizing antibodies in serum (Yu et al., 2020; Mercado et al., 2020; McMahan et al., 2020). These data corroborate previously reported findings on SARS-CoV, showing that neutralizing antibody responses against the SARS-CoV spike protein, that binds to the same cellular receptor as SARS-CoV-2 for cell entry (Shan et al., 2020), were associated with protection against SARS-CoV challenge in nonclinical models (Chen et al., 2005).

Interim analyses of a Phase 1/2a study showed that Ad26.COV2.S elicits a prompt and robust immune response after a single-dose vaccination in both adults (18-55 years old) and elderly (≥65 years old) humans, as measured up to day 29 post-immunization (Sadoff et al., 2021). Based on these data, the protective efficacy against COVID-19 is currently being evaluated in humans in a Phase 3 one-dose efficacy trial (ENSEMBLE trial, NCT04505722). A second Phase 3 study (ENSEMBLE 2, NCT04614948) is currently evaluating vaccine efficacy and durability of two-doses Ad26.COV2.S regimen as well, as the durability of immunity and efficacy may potentially be enhanced by a second dose. Indeed, in other programs with Ad26-based vaccines, two doses induced higher and more durable immune responses (Geisbert et al., 2011; Callendret et al., 2018; Salisch et al., 2019; Salisch et al., 2021). Here we report immunogenicity data after one- and two-dose regimens of Ad26.COV2.S in adult NHP, including a group of aged NHP, for a follow-up period up to 14 weeks after the first vaccination, as well as protective efficacy data in aged NHP challenged with SARS-CoV-2 carrying a glycine residue in position 614 of the spike protein (D614G mutation), which emerged as the most prevalent SARS-CoV-2 spike variant (G614) in the global pandemic thus far.

## Results

### Immunogenicity of one- and two-dose Ad26.COV2.S vaccine regimes in adult rhesus macaques

Adult Rhesus macaques (*Macaca mulatta*; 57 females and 3 males, 3.3 - 5.0 years old) were immunized with either a single dose of 1×10^11^ viral particles(vp) or 5×10^10^ vp Ad26.COV2.S (n=14 per group) or with two doses of 5×10^10^ vp Ad26.COV2.S with a 4- or 8-week interval (n=14 per group). A sham control group (n=4) received an injection with saline at week 0 and week 8. SARS-CoV-2 spike protein-specific antibody responses were measured every two weeks up to 14 weeks after the first immunization by enzyme-linked immunosorbent assay (ELISA) and pseudovirus neutralization assay (psVNA). Immune responses were detected in all vaccinated animals as early as two weeks after immunization and significantly increased by week 4 post-immunization (p≤0.010, ANOVA paired t-test) (Figure 1A and B). Animals that received 1×10^11^ vp Ad26.COV2.S had 1.6-fold higher binding- and 2.1-fold higher neutralizing antibody levels (p=0.008 and p=0.004, respectively, ANOVA t-test) relative to animals immunized with 5×10^10^ vp Ad26.COV2.S. Similar differences in response levels were maintained throughout the entire observation period. However, at week 14, neutralizing antibody titers were similar between the two one-dose groups (p=0.096, ANOVA paired t-test). Spike protein-specific binding antibody levels declined more rapidly than neutralizing antibody levels, irrespective of the vaccine dose the animals had received.

**Figure 1.**
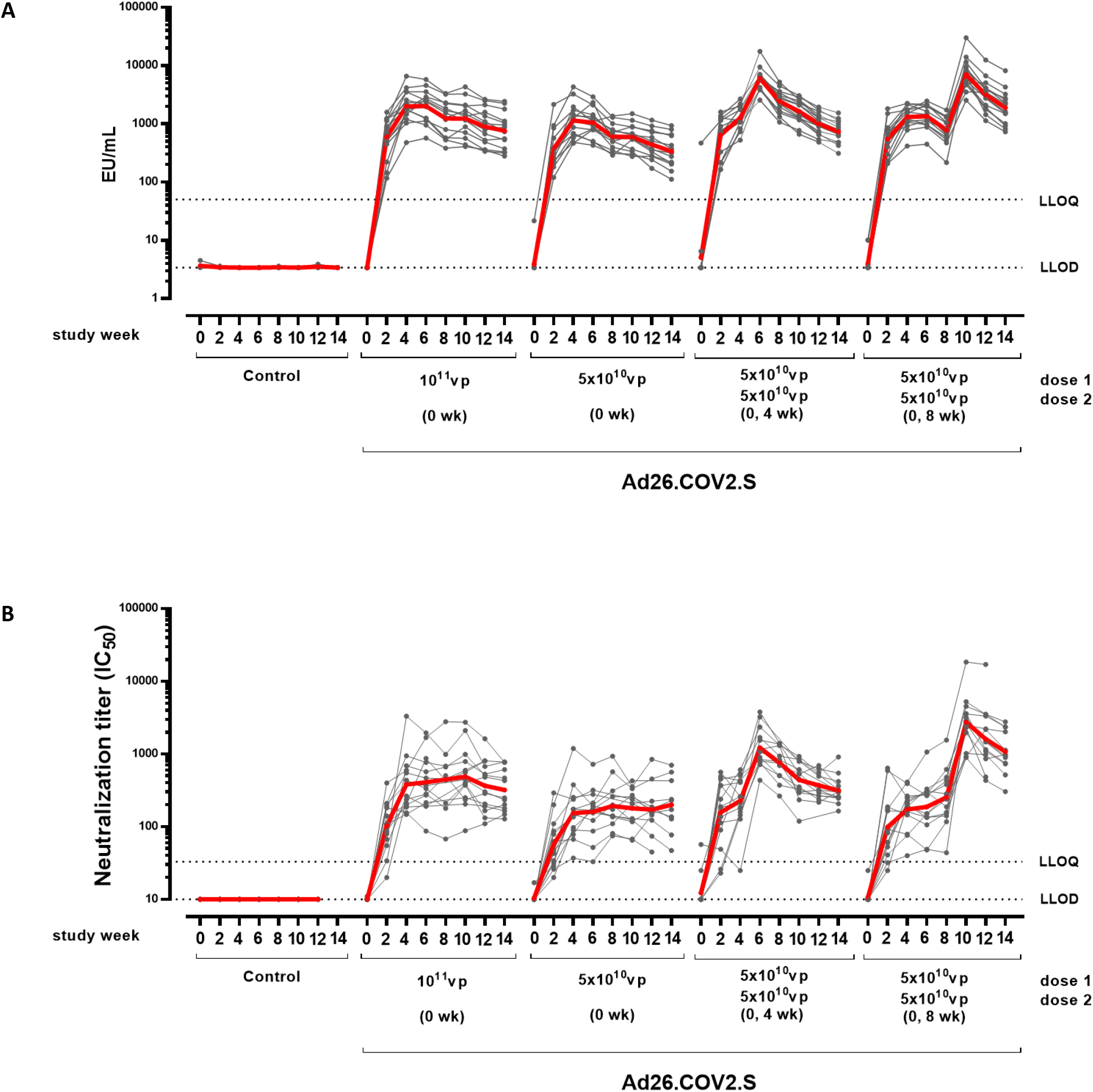
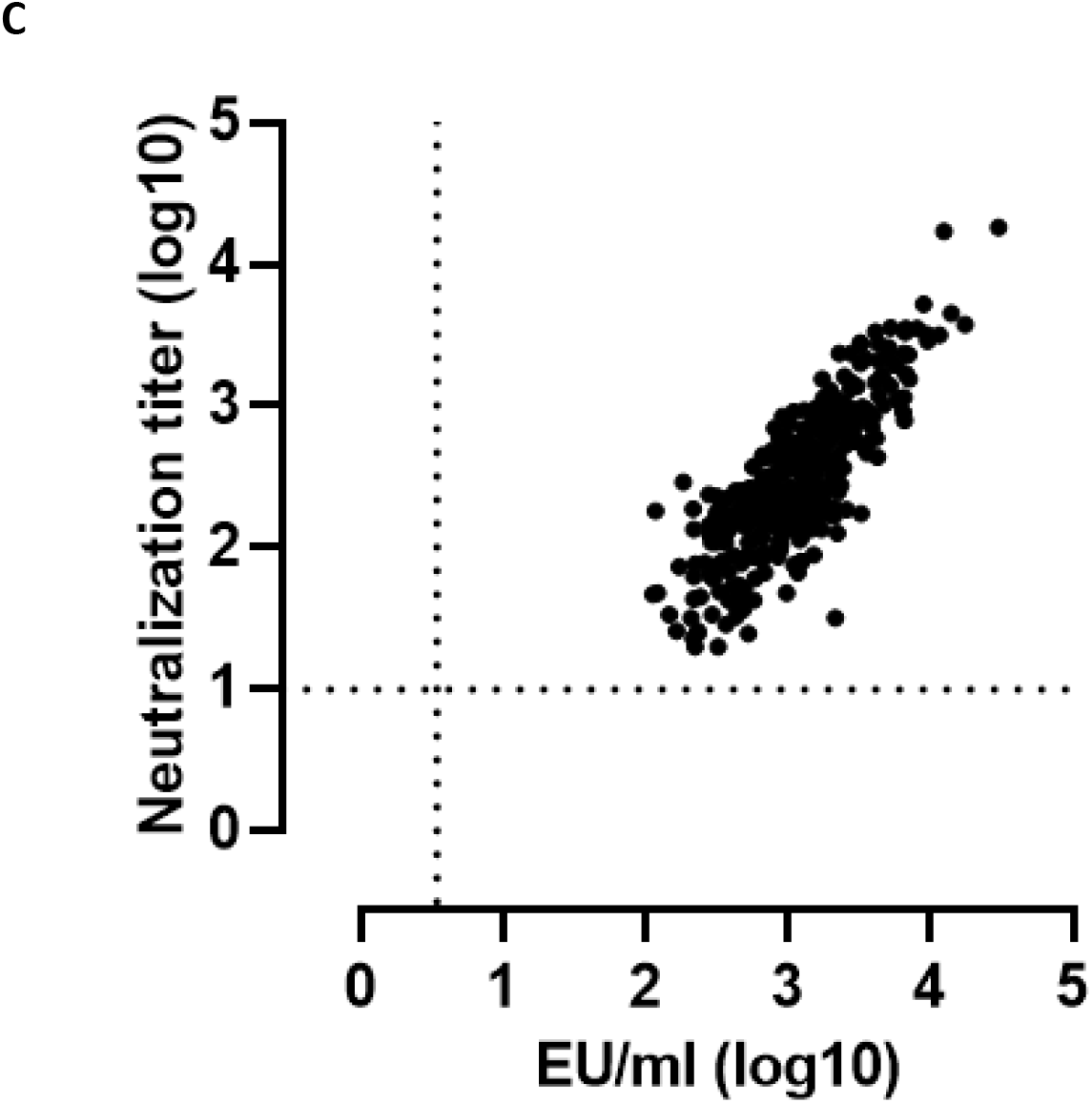
SARS-CoV-2-specific humoral immune responses to one- and two-dose Ad26.COV2.S vaccine regimes in adult rhesus macaques. **A)** SARS-CoV-2 spike protein binding antibody concentrations were measured over time with an ELISA qualified for human samples, using a trimeric, soluble stabilized spike protein produced in mammalian cells as coating antigen. Antibody levels in the individual animals are depicted with grey points and paired measurements connected with grey lines. The geometric mean titers (GMT) of binding antibody responses per group is indicated with the red line. The dotted lines indicate the lower limit of detection (LLOD) and lower limit of quantification (LLOQ). EU/mL; ELISA Units per mL **B)** SARS-CoV-2 spike protein neutralizing antibody titers were measured over time with a psVNA qualified for human samples, using pseudotyped virus particles made from a modified Vesicular Stomatitis Virus (VSVΔG) backbone and bearing the S glycoprotein of SARS-CoV-2. Neutralizing antibody responses are measured as the reciprocal of the sample dilution where 50% neutralization is achieved (IC_50_). Antibody levels in the individual animals are depicted with grey points and paired measurements connected with grey lines. The GMT of neutralizing antibody responses per group is indicated with the red line. The dotted lines indicate the LLOD and LLOQ. **C)** Correlation between spike protein-specific binding antibody concentrations and neutralizing antibody titers per animal for all groups and timepoints except the sham control group and week 0 (baseline). The dotted lines indicate the LLOD for each assay.

A second vaccine dose given 4 or 8 weeks after the first vaccination elicited a significant increase in spike protein-specific antibody responses relative to the pre-dose 2 timepoint (p≤0.001, ANOVA t-test) (Figure 1A and 1B). Compared to the one-dose regimen with 5×10^10^ vp Ad26.COV2.S, a second immunization given at 4 or 8 weeks post first dose, elicited a 5.7- and 11.8-fold increase (p<0.001, ANOVA t-test) in binding antibody concentrations, and a 7.6- and 15.2-fold increase (p<0.001, ANOVA t-test) in neutralizing antibody titers, respectively, as measured 2 weeks post-dose 2. Similar differences were observed when comparing the antibody responses elicited by the two-dose 5×10^10^ vp vaccine regimens, to those elicited by the one-dose 1 x 10^11^ vp vaccine dose.

While the two-dose vaccine regimens with 4- and 8-week interval elicited comparable spike protein-specific binding antibody concentrations two weeks post second immunization (p=0.456, ANOVA t-test) (Figure 1A), the geometric mean of neutralizing antibody titers was 2.2-fold higher for the 8-week compared to the 4-week regimen (p=0.005, ANOVA with t-test) (Figure 1 B). At week 4 and week 6 post second immunization, binding and neutralizing antibody levels declined in both two-dose groups with similar kinetics, maintaining the relative difference in neutralizing antibody titers (2.1- and 2.4-fold higher for the 8-week regimen at 4-and 6-weeks respectively, p=0.021 and p=0.001, respectively, ANOVA t-test).

In spite of the more rapid decline of binding antibody concentrations relative to neutralizing antibody titers in animals that received a one-dose regimen, we observed good overall correlation between binding and neutralizing antibody levels across timepoints for all tested regimens (R =0.7875, p<0.001, Spearman rank-correlation test) (Figure 1C).

### Immunogenicity of one- and two-dose Ad26.COV2.S vaccine regimens in aged rhesus macaques

As COVID-19 severity and mortality increases with age, we additionally analyzed the immunogenicity of Ad26.COV2.S in aged Rhesus macaques (*Macaca mulatta*; 20 females, 13.75 - 21.9 years old). An aluminum hydroxide (Al(OH)_3_) adjuvanted soluble trimeric spike protein stabilized in its prefusion conformation was included as a T helper 2 (Th2) skewing control vaccine for immunological assessment only. Groups were immunized with a one-dose regimen of 1×10^11^ vp Ad26.COV2.S (n=6), a two-dose regimen of 5×10^10^ vp Ad26.COV2.S (n=6) or a two-dose regimen of Al(OH)_3_-adjuvanted 100 μg spike protein (n=4), 8 weeks apart. A sham control group received an Ad26 vector encoding an irrelevant antigen (Ad26.RSV.gLuc; sham control; n=4) at week 0 and week 8. SARS-CoV-2 spike protein-specific binding and neutralizing antibody levels were measured every two weeks up to 10 weeks post the first immunization and spike protein-specific cellular responses were measured at 4 and 10 weeks.

Spike protein-specific binding antibody concentrations significantly increased for each vaccination regimen from week 2 onwards (p≤0.034, ANOVA paired t-test comparing week 0 versus week 2). At weeks 6 and 8 the Ad26.COV2.S induced antibody concentrations were significantly increased compared to Al(OH)_3_-adjuvanted spike protein induced concentrations (p≤0.036, ANOVA t-test). No statistically significant differences in antibody responses elicited by the two regimens employing different Ad26.COV2.S dose levels could be detected up to week 8.

At week 10, two weeks after the second dose, the groups that received a second dose of 5×10^10^ vp Ad26.COV2.S or Al(OH)_3_-adjuvanted spike protein had significantly higher antibody concentrations compared to recipients of the single dose 1×10^11^ vp Ad26.COV2.S (4.4-fold and 5.9 fold for the 5×10^10^ vp Ad26.COV2.S group and Al(OH)_3_-adjuvanted spike protein group respectively, p≤0.002, ANOVA t-test). Spike-specific antibody concentrations between the two dose regimens were not significantly different (p=0.482) (Figure 2A).

**Figure 2.**
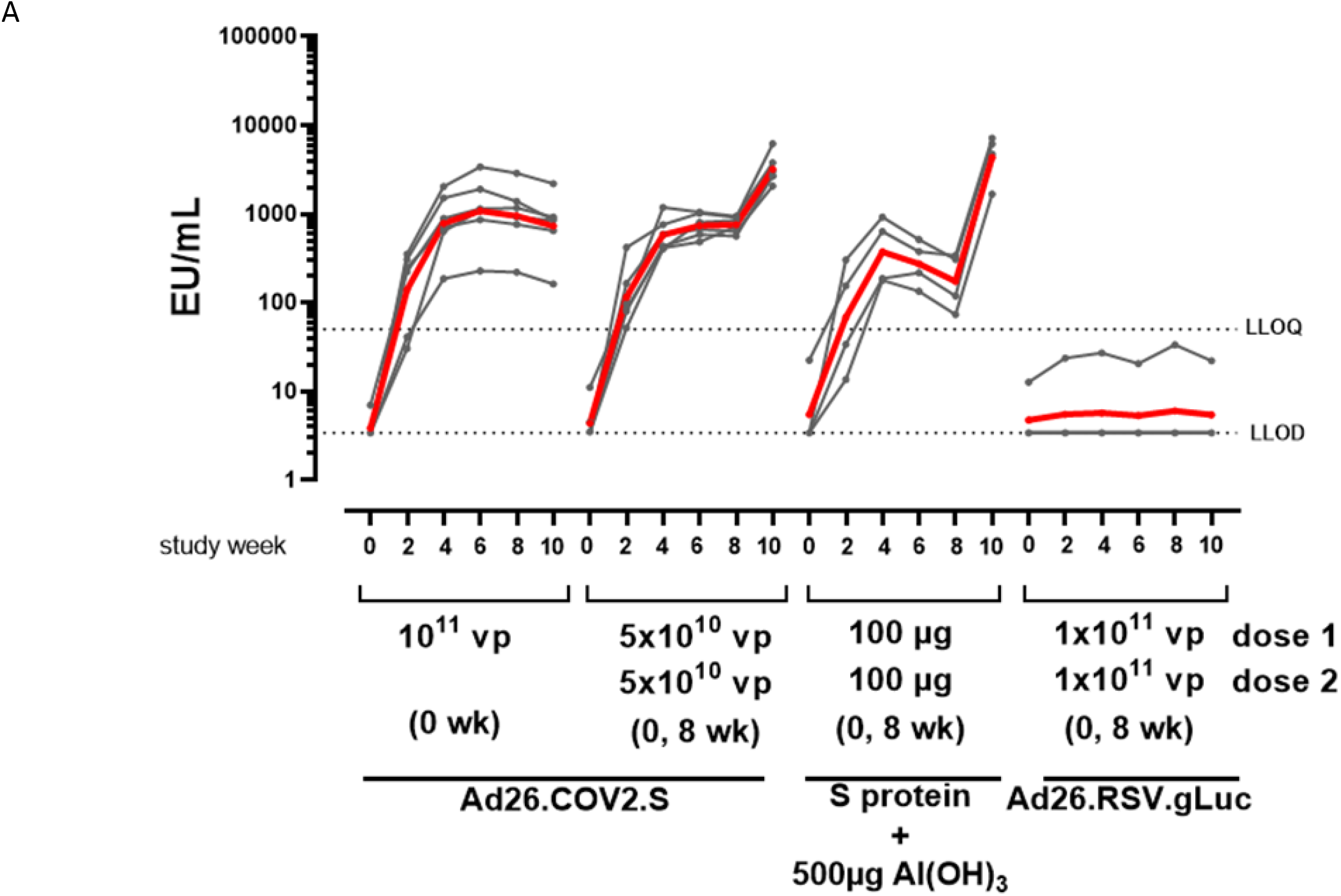
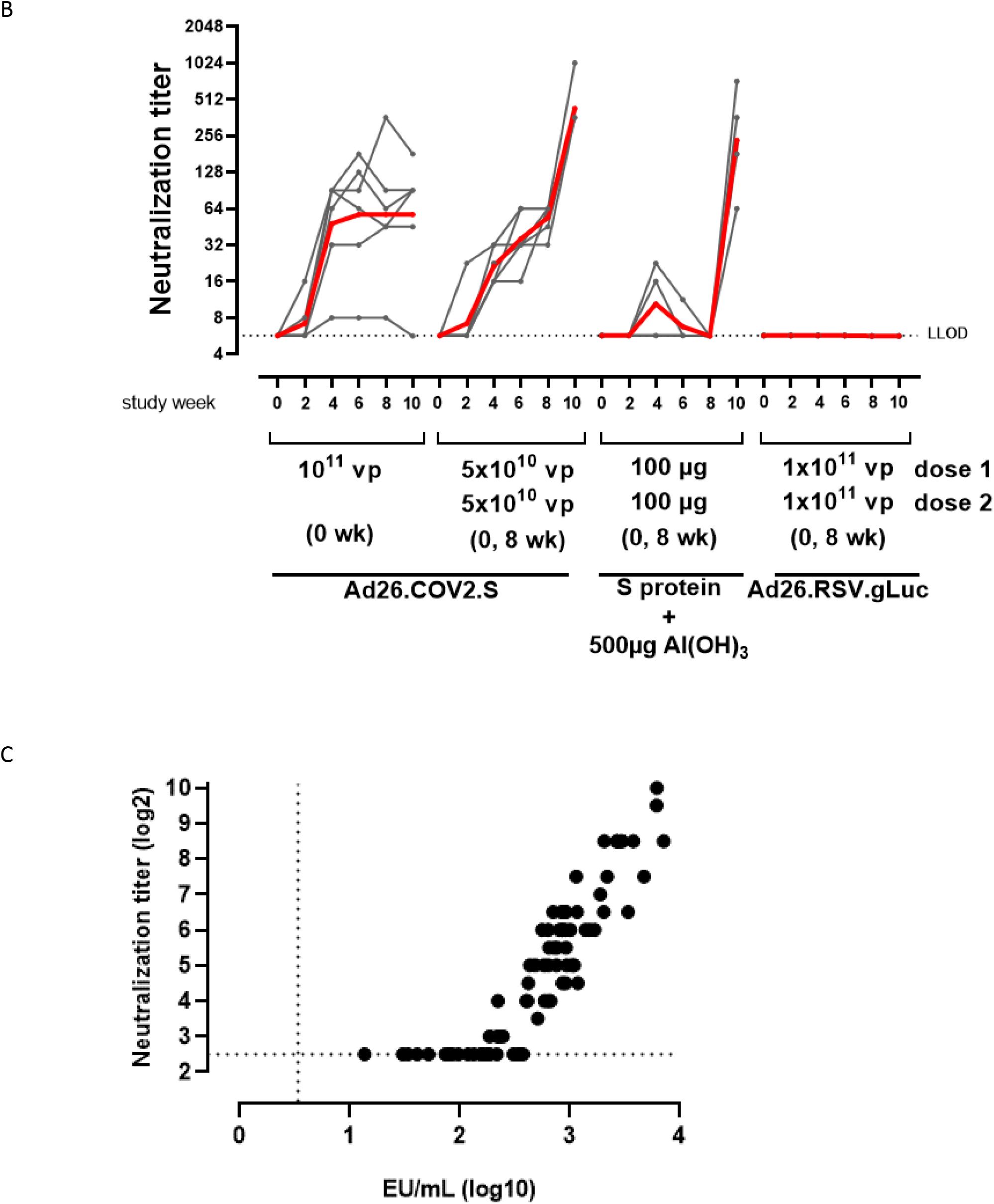
Humoral responses of one- and two-dose Ad26.COV2.S vaccine regimens in aged rhesus macaques. **A)** SARS-CoV-2 spike protein binding antibody concentrations were measured over time with an ELISA qualified for human samples, using a trimeric, soluble stabilized spike protein produced in mammalian cells as coating antigen. Antibody levels in the individual animals are depicted with grey points and paired measurements connected with grey lines. The geometric mean titer (GMT) of binding antibody responses per group is indicated with the red line. The dotted lines indicate the lower limit of detection (LLOD) and lower limit of quantification (LLOQ). **B)** SARS-CoV-2 neutralization antibody titers over time, as measured by wtVNA using the Leiden-0008 strain. Antibody levels in the individual animals are depicted with grey points and paired measurements connected with grey lines. The GMT per group is indicated with the red line. The dotted line indicates the LLOD. **C)** Correlation between spike-specific binding antibody concentrations and neutralizing antibody titers per animal for all groups and timepoints except the sham control group and week 0. The dotted lines indicate the LLOD for each assay.

Kinetics of neutralizing antibody responses were determined by a wildtype Virus Neutralization Assay (wtVNA). A single dose of 1×10^11^ vp Ad26.COV2.S induced neutralizing antibody titers at week 2 which were significantly increased at week 4 in all animals compared to the previous timepoint (p=0.031, sign test), and remained stable thereafter up to week 10. Similarly, the two-dose 5×10^10^ vp Ad26.COV2.S regimen induced neutralizing antibody titers that significantly increased at week 4 (p=0.031, sign test) and 6 (p=0.008, Tobit ANOVA z-test) compared to previous time points. At week 10, 2 weeks after the second dose, antibody titers were increased 8-fold compared to week 8 (p<0.001, Tobit ANOVA z-test). Al(OH)_3_-adjuvanted spike protein induced only low and transient levels of neutralizing antibodies after the first dose in 2 out of 4 animals only. At week 10 however, 2 weeks after the second dose, all 4 animals had neutralizing antibody titers in the same range as the Ad26.COV2.S groups (no statistical analysis possible due to small group size of the adjuvanted protein group). Pairwise comparison of vaccine groups at week 10 showed that the two-dose 5×10^10^ vp Ad26.COV2.S regimen or Al(OH)_3_-adjuvanted spike protein induced significantly higher neutralizing antibody titers compared to the single-dose 1×10^11^ vp Ad26.COV2.S group (10- and 5.5-fold for 5×10^10^ vp Ad26.COV2.S and Al(OH)_3_-adjuvanted spike protein group, respectively, p≤0.004, Tobit ANOVA z-test). Neutralizing antibody titers against the SARS-CoV-2 isolate Leiden-0008, which contains the D614G spike protein mutation (Plante et al., 2020), were not significantly different (p=0.303, Tobit ANOVA z-test) between the two-dose regimens at week 10 (Figure 2B). The spike protein-specific neutralizing antibody titers strongly correlated with binding antibody concentrations (R=0.92, p=<0.001, Spearman rank correlation), showing a higher sensitivity of the ELISA (Figure 2C). There was also a strong correlation observed between neutralizing antibody titers against SARS-CoV-2 isolate Leiden-0008 and the Victoria/1/2020 (D614 variant) in an additional neutralization assay (R=0.89, p=<0.001, Spearman rank correlation; Supplementary figure 1).

Spike protein-specific T cell responses were measured with enzyme-linked immunospot assay (ELISpot) and intracellular cytokine staining (ICS) using peripheral blood mononuclear cells (PBMC) stimulated with 15-mer peptides overlapping by 11 amino acids and spanning the complete SARS-CoV-2 spike protein. Both Ad26.COV2.S regimens as well as Al(OH)_3_-adjuvanted spike protein induced interferon gamma (IFN-γ) responses as measured by ELISpot at 4 weeks after the first dose. At week 10, IFN-γ responses were lower for the 1×10^11^ vp Ad26.COV2.S and adjuvanted spike protein groups compared to week 4. In animals vaccinated with the two-dose 5×10^10^ vp Ad26.COV2.S regimen IFN-γ responses at week 10 were comparable to week 4, suggesting that a second dose of Ad26.COV2.S maintains spike-specific T cell responses. Substantial IL-4 responses were observed only for the Al(OH)_3_-adjuvanted spike protein group at both week 4 and week 10 by ELISpot (Figure 3A).

**Figure 3.**
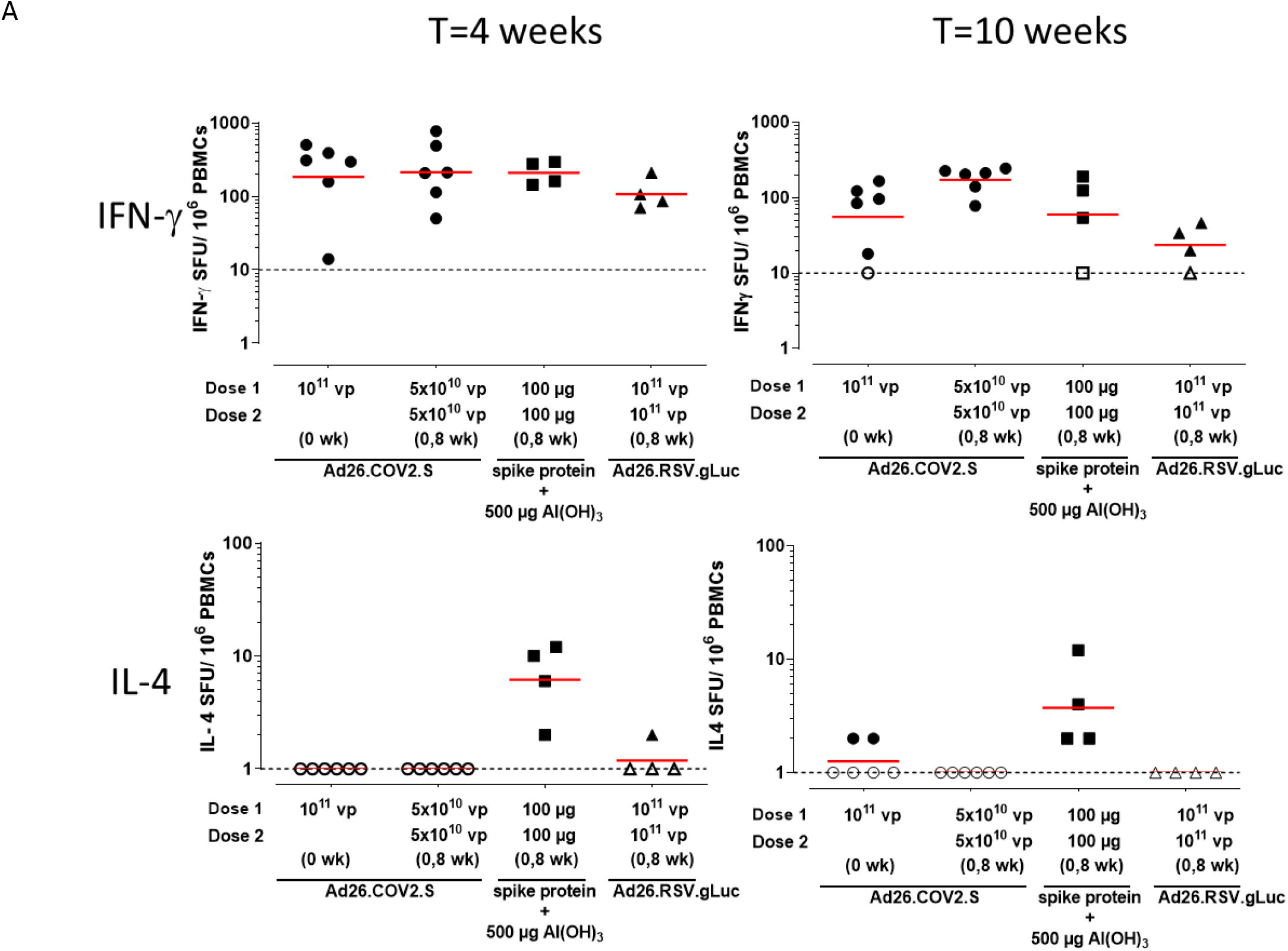
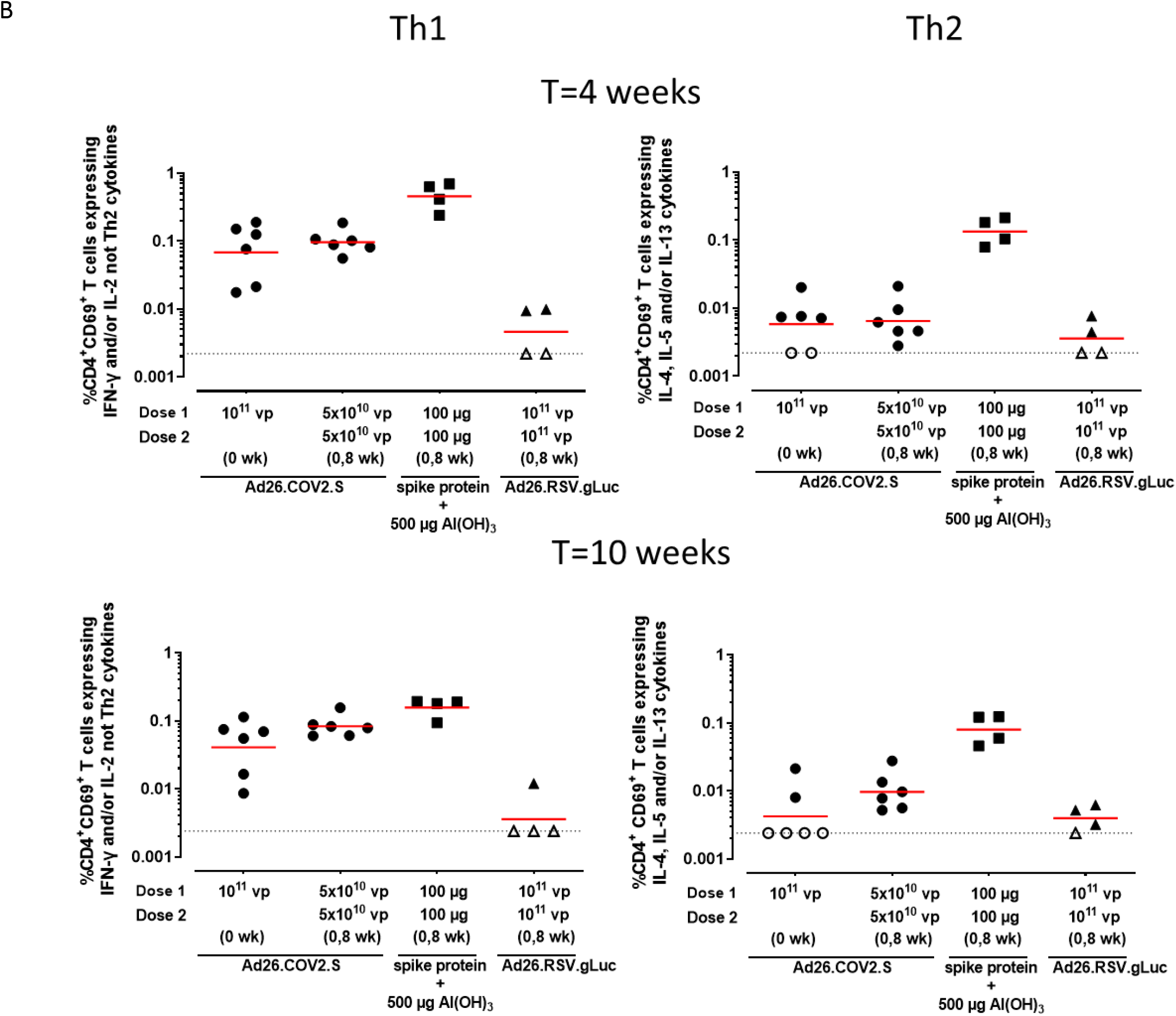
SARS-CoV-2-specific cellular immune responses after vaccination of aged rhesus macaques. **A)** Spike protein-specific T cell responses as measured with an IFN-γ/ IL-4 Double-color ELISpot at indicated timepoints. The geometric mean titer (GMT) response per group is indicated with a horizontal line. Samples with background subtracted counts below or equal to 0 were set at 10 and 1 for IFN-γ and IL-4 respectively for visualization purposes and indicated by open symbols and the dotted line. **B)** Spike protein-specific Th1 and Th2 T cell responses as measured by intracellular cytokine staining at indicated timepoints. Frequency of CD4^+^CD69^+^ T cell expressing Th1 cytokines (IFN-γ and/or IL-2, and not IL-4, IL-5 and IL-13) or Th2 cytokines (IL-4 and/or IL-5 and/or IL-13). Gating strategy is provided in supplemental figure 4. The geometric mean response per group is indicated with a horizontal line. The dotted line indicates the technical threshold. Open symbols denote samples at technical threshold.

CD4+ and CD8+ T cell cytokine responses were also analyzed by intracellular cytokine staining (ICS). Ad26.COV2.S induced a CD4+ Th1-biased response with minimal expression of Th2 cytokines, while Al(OH)_3_-adjuvanted spike protein induced a more dominant Th2 response (Figure 3B and Supplementary figure 2). Spike protein-specific CD8+ T cells induced by Ad26.COV2.S mainly produced IFN-γ and IL-2, while CD8+ T cells induced by Al(OH)_3_-adjuvanted spike protein only produced IL-2. None of the immunization regimens induced CD8+ T cells producing significant amounts of IL-4, IL-5 or IL-13 (Supplementary figure 3).

### Protective efficacy of one- and two-dose Ad26.COV2.S vaccine regimes in aged rhesus macaques

Thirteen weeks after the first Ad26.COV2.S dose, the one-dose 1×10^11^ vp Ad26.COV2.S group, two-dose 5×10^10^ vp Ad26.COV2.S group and the sham control group were inoculated with a total dose of 1×10^5^ tissue culture infective dose 50 (TCID_50_) SARS-COV-2 strain Leiden-0008, by the intranasal and intratracheal route. To increase statistical power, data from a challenge of naïve animals (n=4) using an identical challenge strain, challenge regimen and readouts were added to the sham control group data, collectively referred to as pooled control. Viral loads were assessed by reverse transcription-quantitative polymerase chain reaction (RT-qPCR), measuring subgenomic messenger RNA (sgmRNA) (Wölfel et al., 2020) levels in nasal and tracheal swabs daily during the follow-up period. Low levels of virus were detected in the nose and trachea of some vaccinated animals. In the nose, the median number of days that virus was present in each animal was 0.5 days (range 0-3 days) and in the trachea 2.5 days (range 1-5 days) for the single dose 1×10^11^ vp Ad26.COV2.S group. For the two-dose 5×10^10^ vp Ad26.COV2.S group, median number of days virus was present in the nose of each animal was 1 day (range 0-2 days), and in the trachea 3 days (range 2-4 days). By contrast, in the pooled control group virus was present in the nose for the entire follow-up period for all, except one animal that was consistently negative for nose viral sgmRNA (median of 7 days, range 0-7 days), while in the trachea the median number of days virus was present was 6 days (range 4-7 days) for each animal (Supplementary figure 5). Quantification of total viral load in the follow-up period per animal determined by calculating area under the curve (AUC), showed that total viral load was significantly lower in both vaccinated groups compared to the pooled control group in both samples, from the nose (p≤0.012, Tobit ANOVA z test) as well as from the trachea (p≤0.013, Tobit ANOVA z test; Figure 4A). Bronchoalveolar lavage (BAL) samples were collected at regular intervals during the follow-up period for assessment of viral sgmRNA as well. Only one animal had detectable viral sgmRNA just above the limit of detection in the two-dose Ad26.COV2.S vaccine group, while it was consistently present at high levels in BAL samples from all control animals (Figure 4B and Supplementary figure 5). The single dose Ad26.COV2.S group was not sampled for BAL due to restrictions in the number of animals that can be handled in the Biosafety Level 3 (BSL3) facility. At day 7 and 8 all animals were euthanized and lung tissue was collected. The majority of both one and two-dose Ad26.COV2.S vaccinated animals did not show virus above the limit of quantification in any of the lung lobes tested, while all lung lobes from pooled control animals except one contained viral sgmRNA, with the majority of these at high levels (Figure 4C).

**Figure 4.**
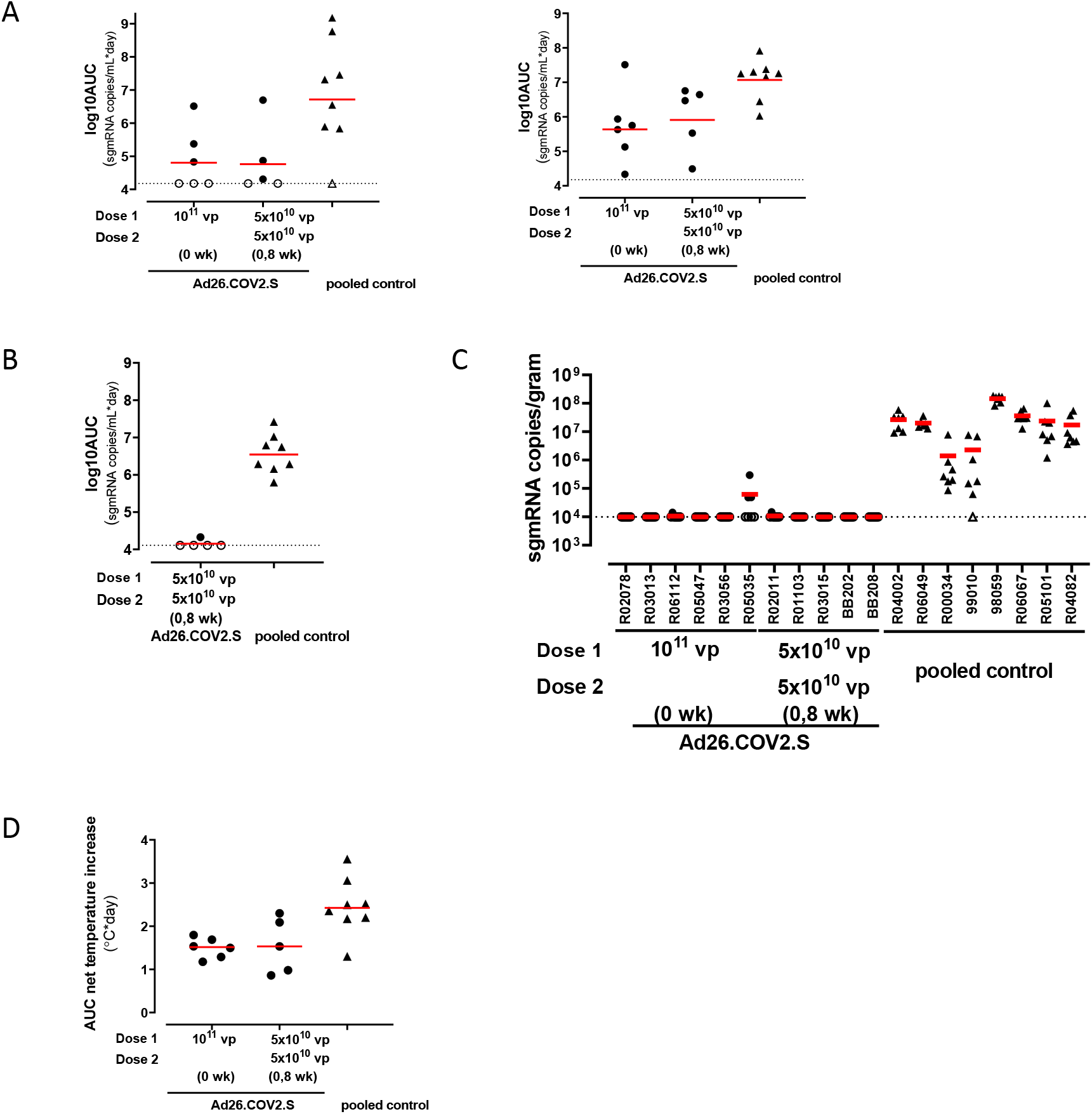
Protective efficacy against SARS-CoV-2 inoculation after vaccination of aged rhesus macaques. Animals were challenged with 1×10^5^ TCID_50_ SARS-CoV-2 administered intranasally and intratracheally Thirteen weeks after the first vaccine dose. Data from a challenge of naïve animals (n=4) using an identical challenge strain, challenge regimen and readouts were added to the sham control group data, collectively referred to as pooled control, to increase statistical power. **A)** Cumulative viral load (sgmRNA) in daily nasal (*left panel*) and tracheal (*right panel*) swabs, defined by area under the curve (AUC) calculation and expressed as log10 AUC (sgm RNA copies/mL x days). Note that for AUC calculation the day of death of all animals was aligned to day 7 to allow combining data from animals euthanized at day 7 and day 8. **B)** Cumulative viral load (sgmRNA) in BAL, obtained every other day during the follow-up period, defined by AUC calculation and expressed as log10 AUC (sgm RNA copies/mL x days). Note that it was only possible to perform BAL on a limited number of animals and it was decided to exclude the one-dose 1×10^11^ vp Ad26.CoV2.S group. **C)** Viral load (sgmRNA) in lung tissue. Viral load was measured of each individual lung lobe (7) of each animal and expressed as log10 sgm RNA copies/gram. A lower right lung lobe sample of one animal in the pooled control group was not available. **D)** Fever duration, defined as AUC of the net temperature increase during the first 6 consecutive days of the follow-up period relative to a pre-challenge baseline period. Red horizontal lines represent group geometric mean titers, the dashed horizontal line indicates the lower limit of quantification (LLOQ). Open symbols denote samples at LLOQ in all panels.

The animals showed no overt clinical signs after virus infection and clinical chemistry parameters in blood were normal Body temperature was continuously measured throughout the study. Pre-challenge data were used to reconstruct a daily baseline temperature profile for each individual animal. Fever, defined as temperature increase above baseline was recorded post-infection and total temperature increase during the follow-up period was calculated by means of AUC (fever duration). All animals showed an increase in temperature after SARS-CoV-2 inoculation. A modest yet statistically significant reduction in fever duration was observed for both vaccinated groups compared to the pooled control animal group (p≤0.012, t-test; Figure 4D and Supplementary figure 6A-C).

Histological analysis of lung tissue at the end of the study showed minimal pulmonary pathology in vaccinated animals, with minimal to mild mononuclear (macrophages and lymphocytes) or mixed cell infiltrates (macrophages, lymphocytes and scattered neutrophils). in the interstitium and alveolar lumina minimal to mild perivascular cuffing and focal bronchiolo-alveolar hyperplasia was observed (Figure 5A -B -D, -E). Immunohistochemistry staining for SARS-CoV-2 nucleoprotein (N) detected a few isolated positive pneumocytes in a single lung lobe in 1 out of 11 vaccinated animals. (Figure 5 G-H). By contrast, sham control animals demonstrated evidence of viral interstitial pneumonia, characterized mostly by moderate mononuclear or mixed cell infiltrate in the interstitium, mild to moderate type II pneumocyte hyperplasia/bronchiolo-alveolar hyperplasia and alveolar lumina containing edema (homogenous eosinophilic fluid) admixed with mononuclear or mixed cell infiltrates (Figure 5C, -F). In 7 out of 8 sham Ad26.RSV.gLuc vaccinated sham control animals, minimal to moderate numbers of SARS-CoV-2 N-positive pneumocytes were noted in multiple lung lobes (Figure 5I) consistent with RT-qPCR data from the lung lobes.

**Figure 5.**
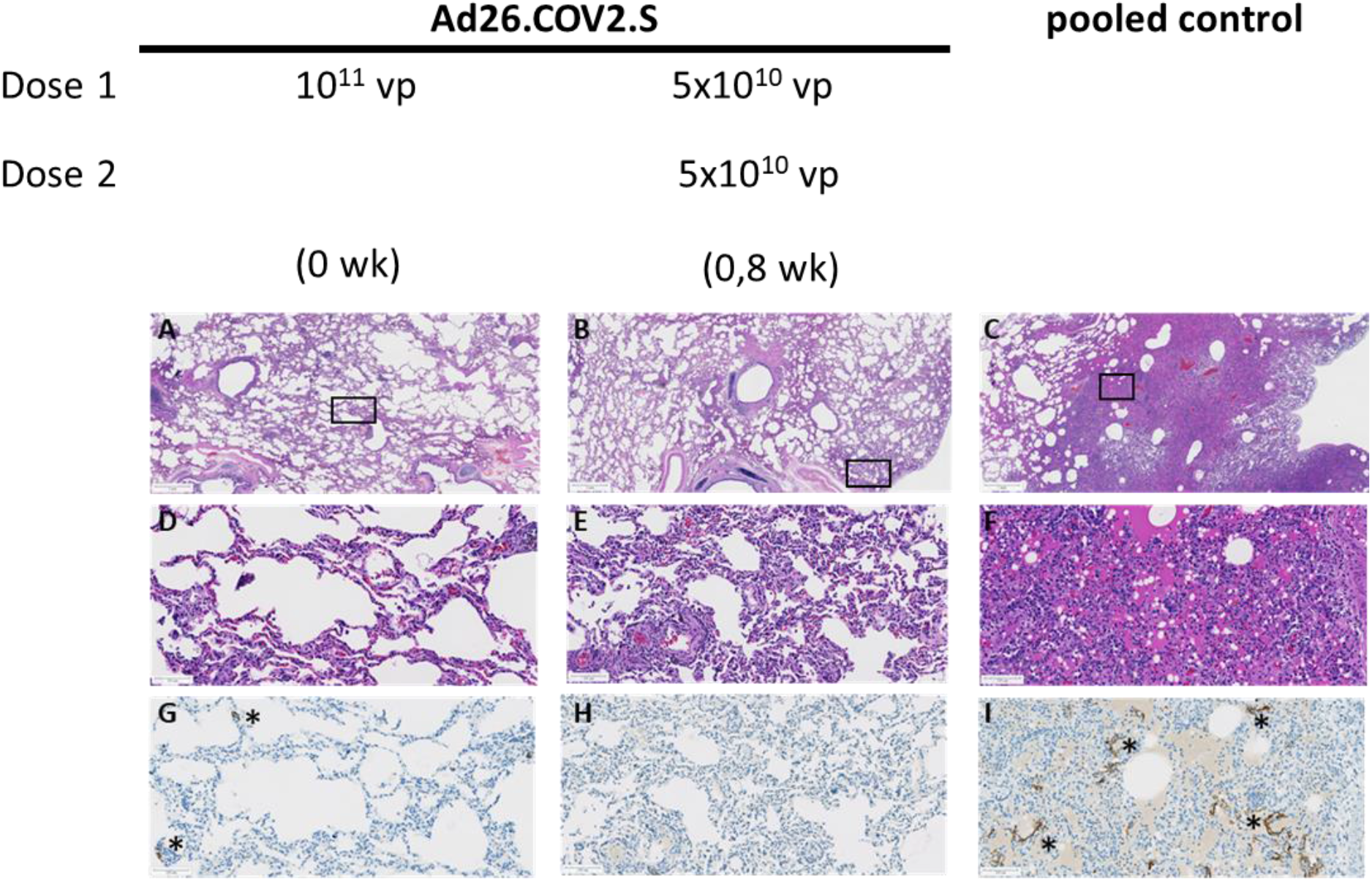
Histology after SARS-COV-2 inoculation of vaccinated aged rhesus macaques, Seven individual lung lobes were evaluated for each animal .in each treatment group **A-C)** 10x magnification overview image per treatment group. Scale bar is 1 mm. **D-F)** hematoxylin/eosin staining of rectangular areas as indicated in panels A-C, from an Ad26.RSV.gLuc-vaccinated animal. **D)** (*). In the 1×10^11^ vp Ad26.COV2.S group minimal mononuclear cell interstitial infiltrate and minimal perivascular cuffing was observed. **E)** In the 5×10^10^ vp Ad26.COV2.S group multifocal small areas with minimal mixed-cell interstitial infiltrates (macrophages, scattered neutrophils and lymphocytes) and minimal perivascular cuffing were observed. Alveolar lumina contained minimal macrophages, lymphocytes and scattered neutrophils. **F)** In the pooled control group animals, here represented by an animal of the Ad26.RSV.gLuc sham control group, focally extensive to diffuse lesions were observed. with moderate mononuclear cell infiltrate in interstitium (macrophages, lymphocytes) and mild type II pneumocyte hyperplasia. Alveolar lumina contained edema, macrophages, lymphocytes and scattered neutrophils. **G-I)** SARS-CoV-2 -N immunohistochemistry (red-brown staining) of rectangular areas as indicated in panels A-C. Antigen-positive pneumocytes are marked with an asterisk (*). In the 1×10^11^ vp Ad26.COV2.S group, one out of 6 animals had a single lung lobe were staining was observed. G) Focal area with individual SARS-CoV-2 N positive pneumocytes (*) in the left caudal lung lobe H) Absence of SARS-CoV-2 N positive pneumocytes in all lung lobes from all animals in the 5×10^10^ vp Ad26.COV2.S group. I) Multifocal minimal to moderate numbers of SARS-CoV-2 N positive pneumocytes (*) in 7 out of 8 pooled control animals. Panels D-I are 100x magnification, scale bar is 100μm.

## Discussion

We previously reported immunogenicity and protective efficacy data of a single dose of our COVID-19 vaccine candidate Ad26.COV2.S in adult NHP (Mercado et al., 2020). Here we evaluated the immunogenicity of one- and two-dose Ad26.COV2.S regimens in adult and aged rhesus macaques for up to 14 weeks after the first dose, to gain insight into the durability of immunity after a single dose of this vaccine candidate and the added value of a second dose on the magnitude of spike protein-specific immune responses. In addition, we assessed the protective efficacy of one- and two-dose Ad26.COV2.S regimens in aged NHP. We used a new challenge model based on the D614G spike SARS-CoV-2 variant, which is since spring 2020 the most prevalent circulating spike variant thus far.

In both adult and aged macaques, spike protein-binding and SARS-CoV-2 neutralizing antibody responses were detected as early as two weeks after the first Ad26.COV2.S immunization and were significantly increased by week 4 in complete agreement with our clinical trial observations in adults and elderly after a single immunization with Ad26.COV2.S (Sadoff et al., 2021)

Humoral immune responses were maintained at least up to week 14 post-immunization, providing an early sign of the durability of immunity elicited by Ad26.COV2.S. Binding and neutralizing antibody responses showed some decline over time, while neutralizing antibody responses appeared to be more stably maintained, especially in recipients of the 5×10^10^ vp dose level. Although humoral immune responses were initially significantly higher in NHP that received the 1×10^11^ vp dose as compared to recipients of the 5×10^10^ vp vaccine dose, differences in neutralizing antibody levels decreased over time and do not suggest a clear benefit of the higher dose, in agreement with interim Phase 1/2a clinical data (Sadoff et al., 2021).

A second dose of Ad26.COV2.S given with an 8 week interval resulted in a significant increase in spike protein-specific binding and neutralizing antibody responses, in both adult and aged NHP. This is in line with our data in humans (Sadoff et al., 2021) and with observations with other Ad26-based vaccines (Geisbert et al., 2011; Callendret et al., 2018; Salisch et al., 2019; Baden et al., 2013; Salisch et al., 2021).

Neutralizing antibody titers were higher in animals that received the two vaccine doses 8-weeks apart as compared to the 4-week interval, albeit both two-dose regimens were more immunogenic than the one-dose regimen. This confirms that a longer interval between vaccine doses can significantly improve the magnitude and/or quality of the antibody response. (Ledgerwood et al., 2013; Siegrist, 2018; Sallusto et al., 2010; Roozendaal et al., 2020) While we have not evaluated the potential difference in efficacy of longer and shorter two-dose regimens in NHP, a two-doses regimen of 5×10^10^ vp Ad26.COV2.S administrated 8-weeks apart is currently being tested for efficacy in humans (ENSEMBLE 2, NCT04614948).

Despite the fact that different virus neutralization assays were used, we observed a strong correlation between binding and neutralizing antibody titers in sera from vaccinated adult and aged NHP across time points. These data confirm our earlier observation (Yu et al., 2020; Mercado et al., 2020; McMahan et al., 2020) and suggest that spike protein-binding antibody concentrations measured by ELISA could be used as a surrogate readout for neutralizing antibody responses.

Almost all Ad26.COV2.S vaccinated aged NHP that were challenged three months after first immunization, were protected from lung infection, as demonstrated by negative PCR testing for sgmRNA in BAL and lung tissue samples. Only one Ad26.COV2.S vaccinated animal that received a single vaccination, had clearly detectable SARS-CoV-2 sgmRNA levels as well as traces of viral antigen detected by immunohistochemistry in lung tissue samples. This animal had much lower binding and neutralizing antibody levels after vaccination, which could explain the breakthrough infection, as protection from infection in earlier studies was correlated with binding and neutralizing antibody titers (Mercado et al., 2020; Yu et al., 2020; McMahan et al., 2020) Nevertheless, viral load in this animal was lower as compared to viral load in lungs of challenged control animals. While complete protection apparently requires a higher neutralizing antibody titers in this particular animal model, it is tempting to speculate that the viral load reduction in lung associated with lower neutralizing antibody titers could translate in protection from severe disease even in low human vaccine responders. Indeed, while histological analysis and immunohistochemistry on lung tissue showed severe pulmonary histopathology and presence of viral antigens in challenged control animals, only minimal histopathological abnormalities and viral antigens in lungs of Ad26.COV2.S vaccinated animals was observed, in agreement with earlier observations (Corbett et al., 2020). We observed a small but persistent decrease in the febrile response post-infection in vaccinated animals. Although transient fever has been described after SARS-CoV-2 infection of rhesus macaques (Munster et al., 2020), no effect on temperature was observed in recent NHP vaccine efficacy studies (Chen et al., 2020) (Wang et al., 2020). In addition to differences in respect to the NHP models used in other studies like age and challenge virus, an important advantage in this study might be that the body temperature was continuously monitored and no additional interventions, were required to record temperature. Continuously temperature monitoring might therefore be a useful clinical parameter in NHP vaccine efficacy models.

The only partial protection of the upper respiratory tract observed in our present study seems at odds with our previous NHP study, in which Ad26.COV2.S elicited immunity provided complete and near-complete protection against viremia in lung and upper respiratory tract, respectively (Mercado et al., 2020). Several factors may contribute to this difference in outcome. NHP in our current study were aged and the time of challenge after immunization was considerably longer, albeit that antibody titers were not waning. Additionally, the G614 SARS-CoV-2 challenge strain instead of the Washington D614 challenge strain was used. and was reported to be associated with enhanced viral replication in the upper respiratory tract and potential enhanced viral transmissibility, but with no associated increase in disease severity (Plante et al., 2020; Hou et al., 2020).

It is important to emphasize that Ad26.COV2.S elicited immunity is protective against SARS-CoV-2 with either the D (Mercado et al., 2020) or the G at amino acid position 614 of the spike protein. However, with newly emerging variants, the constant monitoring of strain coverage by vaccine elicited immunity will continue to be of utmost importance.

Derisking for the potential and theoretical risk of Vaccine-Associated Enhanced Respiratory Disease (VAERD)(Lee et al., 2020; Bottazzi et al., 2020; Haynes et al., 2020), which is generally considered to be associated with non-neutralizing antibody responses and Th2-skewed cellular immunity, is an important aspect in the development of COVID-19 vaccines. Here we show that in aged NHP, Ad26.COV2.S elicited CD4+ T cell responses that were Th1 skewed, confirming our observations in elderly humans (Sadoff et al., 2021) and similar to findings with other genetic vaccine platforms encoding SARS-CoV-2 spike protein(van Doremalen et al., 2020; Yu et al., 2020; Anderson et al., 2020; Vogel et al., 2020; Corbett et al., 2020). The ability of NHP to develop a Th2 skewed immune response was demonstrated by vaccination with an Al(OH)_3_-adjuvanted spike protein. The Th1-skewed response in Ad26.COV2.S vaccinated NHP together with the induction of robust and durable neutralizing antibody responses by Ad26.COV2.S and absence of enhanced lung pathology in challenged animals, indicate that the potential for VAERD with this vaccine is extremely unlikely.

Overall, the immunogenicity and protective efficacy data presented in this manuscript further support our decision to evaluate a single 5×10^10^ vp dose of Ad26.COV2.S in our Phase 3 ENSEMBLE (Trial Number: NCT04505722) study and also to evaluate a two-dose Ad26.COV2.S regimen in our second Phase 3 study ENSEMBLE 2 (Trial Number: NCT04614948)..

## Supporting information

Spplemtary Fifure 1 to 6

## Author Contributions

Designed studies and reviewed data: L.S., H.K., S.K.R.H., J.E.M.vd.L., L.D., R.R., F.W., R.C.Z. Performed experiments and analyzed data: D.N.C.C., A.I.G., M.R.M.B., J.D., J.V., E.v.H., Y.C, J.V., S.K., A.H.d.W., E.K., T.J.D., S.K.M., M.K., E.J.S., K.P.B., M.A.S., I.K., E. J. V., B.E.V., G.K., P.M., W.M.J.M.B., M.v.H., L.M., J.T.B.M.T. Drafted the paper: L.S., H. K. Reviewed the paper: all authors.

## Acknowledgements

This project was funded in part by the Department of Health and Human Services Biomedical Advanced Research and Development Authority (BARDA) under contract HHS0100201700018C. We thank Johan Verspuij for assistance with data processing, Sarah Janssen for ad-hoc statistical support, Janssen colleagues of the Vector Generations and Sub-Unit Vaccine Design departments for providing reagents. We are grateful to colleagues of the Non-Clinical Safety Toxicology/Pathology depart of Janssen Research and Development in Beerse, Belgium for performing histology staining and analysis. We thank Gert Scheper, Danielle van Manen Martin Friedrich Ryser and Joanne Wolter for reviewing the paper and for providing valuable input. We thank Daniella Mortier, Gwendoline Kiemenyi-Kayere, Zahra Fagrouch, Nikki van Driel, Ivonne Nieuwenhuis, Herman Oostermeijer, Lisette Meijer, Henk Niphuis, Ed Remarque, Tom G.M. Haaksma and Boudewijn Ouwerling from the Biomedical Primate Research Centre for logistical and biotechnical support. We thank the team of Charles River Laboratories Montreal ULC, Laval Site (CR-LAV) (Canada) and Reno NV Site (US) for their accurate and punctual work on the NHP study with adult macaques, in particular we thank Anne Marie Downey, Roula Salame, Carolyne Dumont, Rajen Patel and Sunjay Sethi. We thank the Nexelis Team of the Laval site, Canada, for their speed and flexibility in accommodating spike protein ELISA and psVNA sample analysis within short timelines, in particular we thank Luc Gagnon, Helen Diamantakis, Mary Osei-Twum, Greg Kulnis, Steven-Phay Tran, Julien St-Jean, Marcel Dupelle and Akeel Baig. We thank the MNA Testing Team, Public Health England, Porton Down, Salisbury, UK, for the immunogenicity testing, in particular we thank Dr Sue Charlton, Dr Lorna McInroy, Anna England, Durga Rajapaksa, Stephanie Leung, Lauren Allen, Emily Brunt, Dr Kevin Bewley, Dr Naomi Coombes, Imam Shaik, and dr Holly Humphries. From Leiden University Medical Center, we thank Ali Tas for his assistance with preparing the SARS-CoV-2/Leiden-0008 challenge virus used in this study and Shessy Torres for excellent technical support.

## Competing interests

The authors declare no competing financial interests. L.S., H.K., S.K.R.H., J.E.M.vd.L., L.D., D.N.C.C, A.I.G, M.R.M.B., J.D., J.V., E.v.H., Y.C., J.V., A.H.d.W., E.K., J.C. M.v.H., L.M., J.T.B.M.T., R.R., J.C., H.S, F.W. and R.C.Z. are employees of Janssen and may be Johnson & Johnson stockholders.

## Materials and methods Animals

### Adult NHP

The NHP study of adult animals was conducted at Charles River Laboratories (CRL) Montreal ULC, Laval Site (CA). Animals were obtained from Kunmings Biomed international Ltd, China. Prior to transfer from test facility colony, all animals were subjected to a health assessment and tested at least once for tuberculosis by intradermal injection of tuberculin. An anthelmintic treatment was administered to each animal by subcutaneous injection. The evaluations were performed in accordance with the standard operating procedures by technical staff. Animal experiment approval was provided by the Institutional Animal Care and Use Committee (IACUC) at CRL Montreal ULC, Laval Site (CA). Animal experiments were performed in compliance with Guidelines published by the Canadian Council on Animal Care and the Guide for the Care and Use of Laboratory Animals published by the National Research Council Canada. The Test Facility is accredited by the Canadian Council on Animal Care (CCAC) and the American Association for Accreditation of Laboratory Animal Care (AAALAC). In addition, the study was conducted according to EMA guideline, ICH M3(R2): Guidance on Non-Clinical Safety Studies for the Conduct of Human Clinical Trials and Marketing Authorization for Pharmaceuticals and FDA guideline, Redbook 2000: General Guidelines for Designing and Conducting Toxicity Studies.

### Aged NHP

The study using aged NHP was performed at the Biomedical Primate Research Centre, Rijswijk, The Netherlands (an AAALAC-accredited institution). Animals were captive-bred for research purposes and socially housed. Animal housing was according to international guidelines for NHP care and use (The European Council Directive 2010/63, and Convention ETS 123, including the revised Appendix A as well the ‘Standard for humane care and use of Laboratory Animals by Foreign institutions’ identification number A5539-01, provided by the Department of Health and Human Services of the United States of America’s National Institutes of Health (NIH)). The study was conducted in compliance with, and approved by, all relevant local and national regulations and the Institutional Animal Welfare Body (Instantie voor Dierenwelzijn, IvD) guarded that all possible precautions were taken to ensure the welfare and to avoid any unnecessary discomfort to the animals.

## Vaccines

The Ad26.COV2.S vaccine has been generated as previously described (Bos et al., 2020). Briefly, Ad26.COV2.S is a replication-incompetent Ad26 vector encoding a prefusion-stabilized SARS-COV-2 spike protein sequence (Wuhan Hu1; GenBank accession number: MN908947). Replication-incompetent, E1/E3-deleted Ad26-vectors were engineered using the AdVac system (Abbink et al., 2007), using a single plasmid technology containing the Ad26 vector genome including a transgene expression cassette. The human codon optimized, prefusion-stabilized, SARS-COV-2 spike protein encoding gene was inserted into the E1-position of the Ad26 vector genome. Manufacturing of the Ad26 vector was performed in the complementing cell line PER.C6 TetR (Wunderlich et al., 2018; Zahn et al., 2012). The negative control vector Ad26.RSV.gLuc encodes the RSV F protein fused to *Gaussia* firefly luciferase as a single transgene separated by a 2A peptide sequence, resulting in expression of both individual proteins. Manufacturing of the vector was performed in PER.C6 (Sanders et al., 2013).

The full-length spike protein used for immunization (COR200099) (Bos et al., 2020) was produced on Expi293F cells. COR200099 is based on the Wuhan-Hu-1 SARS-CoV-2 strain (MN908947) and stabilized by two point mutations (R682A, R685G) in the S1/S2 junction that disrupts the furin cleavage site, and by two consecutive prolines (K986P, V987P) in the hinge region in S2. In addition, the transmembrane and cytoplasmic regions have been replaced by a fibritin foldon domain for trimerization and a C-tag, allowing the protein to be produced and purified as soluble protein. Adenoviral vectors and protein were tested for bioburden and endotoxin levels prior to use.

## Study design animal experiments

### Adult NHP

60 (57 females and 3 males. 3 males were allocated to test groups 3, 4 and 5, 1 male in each group) rhesus macaques (*Macaca mulatta*) from Chinese origin between 3.3 to 5.0 years of age and weighting between 2.9 to 8.1 kg, were assigned to five groups by a randomizing stratification system based on body weights. Fourteen animals were included in each vaccine group and four animals were included in the sham control group. Group 1 (n=4) is the sham control group and received saline injection at week 0 and week 8. group 2 and 3 (n=14 each group) received one immunization with 1×10^11^ vp and 5×10^10^ vp of Ad26. COV2.S, respectively, at week 0. Group 4 and 5 (n=14 each group) received two immunizations with 5×10^10^ vp of Ad26. COV.2 spaced by four (week 0 and week 4) and eight weeks (week 0 and week 8), respectively. All immunizations were performed via the intramuscular route in the quadriceps muscle of the left hind leg. Blood for serum was obtained prior to the first vaccine dose and every 2 weeks subsequently up to week 14 of the study.

### Aged NHP

20 female rhesus macaques (*Macaca Mulatta*), aged between 13.75 and 21.9 years of age and weighting between 6.6-12.6 kg, were distributed over 4 experimental treatment groups and housed in ABSL-III facilities, pair-housed with socially compatible animals. Prior to study start an AnipillV2 telemetry system (BodyCAP, Hérouville Saint Clair, France) was surgically implanted in the abdomen of animals and recorded body temperature every 15 minutes. Group 1 (n=6) received 1×10^11^ vp of Ad26. COV2.S at week 0. Group 2 (n=6) received 5×10^10^ vp of Ad26. COV2.S at week 0 and 8. Group 3 (n=4) received 100 μg spike protein, adjuvanted with 500 μg Aluminum Hydroxide (Al(OH)_3_; 2% Alhydrogel, InvivoGen) at week 0 an 8. The sham control group (Group 4, n=4) was immunized with 1×10^11^ vp Ad26.RSV.gLuc, an Ad26 vector expressing an irrelevant antigen. All immunizations were performed intramuscularly in quadriceps of the left hind leg. Blood for serum and PBMC isolation was obtained as indicated in the text. Five weeks after the second vaccination dose all groups, except the Al(OH)_3_-adjuvanted spike protein group, were inoculated with 1×10^5^ TCID_50_ of SARS-CoV-2 isolate Leiden-0008. Clinical isolate SARS-CoV-2/human/NLD/Leiden-0008/2020 (Leiden-0008) was isolated from a RT-PCR positive throat swab and passaged twice in Vero E6 cells. The spike protein of this isolate contains the D614G mutation. The NGS-derived complete genome sequence of this virus isolate is available under GenBank accession number MT705206.1 and showed only minor variants from the consensus sequence, especially in the spike furin cleavage site region we detected below 2% of heterogeneity. Isolate Leiden-0008 was propagated and titrated in Vero E6 cells. The inoculum was administered in a 2 mL volume, 1 mL intratracheally and 1 mL intranasally, 0.5 mL per nostril. After virus inoculation, nose and trachea swabs were taken daily, as well as BAL every other day from the two-dose 5×10^10^ vp Ad26. COV2.S and sham control groups, to measure viral load. As animals were anaesthetized on a daily basis, tube feeding was applied. Animals were euthanized at day 7 and 8 after virus inoculation, with the number of animals of each group approximately distributed over both days, and respiratory tract tissues were isolated for histopathology, immunohistochemistry and viral load. To increase statistical power, the data from the sham control group was pooled with data from the pilot virus inoculation study, consisting of naïve animals (n=4) of the same age range that were inoculated identically as the described above for the vaccine study. One animal in the 5×10^10^ vp Ad26.COV2.S group died during the study. Postmortem autopsy identified a marked to severe, acute bronchopneumonia associated with foreign particulate material in the airways, which is consistent with aspiration pneumonia in the lung as the cause of death of this animal. The death was therefore deemed unrelated to the vaccine and the animal was excluded from all other analyses.

## Enzyme-linked immunosorbent assay (ELISA)

IgG binding to SARS-CoV-2 spike protein was measured by ELISA using a recombinant spike protein antigen based on the Wuhan-Hu-1 SARS-CoV-2 strain (MN908947). The SARS-CoV-2 spike protein antigen was adsorbed on 96 well microplates for a minimum of 16 hours at 4°C. Following incubation, plates were washed in phosphate-buffered saline (PBS)/0.05% Tween-20 and blocked with 5% skim milk in PBS/0.05% Tween-20 for 1 hour at room temperature. Serum standards, controls and NHP serum samples were diluted and incubated on the plates for 1 hour at room temperature. Next, the plates were washed and incubated with peroxidase conjugated goat anti human IgG for 1 hour at room temperature, washed, and developed with tetramethylbenzidine (TMB) substrate for 30 minutes at room temperature and protected from light, then stopped with H2SO4. The optical density was read at 450/620 nm. The antibody concentrations were back calculated on the standard and the reportable value were generated based on all passing dilutions, expressed in ELISA units [EU]/mL. The LLOD is 3.4 EU/mL, based on the standard lowest interpolation range concentration multiplied per the dilution factor and is used as an informative LLOD. LLOQ is based on qualification performed for human samples and has been set on 50.3 EU/mL.

## Pseudovirus neutralization assay (psVNA)

SARS-CoV-2 S neutralizing antibody titers were measured by pseudovirus neutralizing assay. Pseudotyped virus particles were made from a modified Vesicular Stomatitis Virus (VSVΔG) backbone and bear the S glycoprotein of the SARS-CoV-2. The pseudoparticles contain a luciferase reporter gene used for detection. Serial dilutions of heat-inactivated NHP serum samples were prepared in 96-well transfer plates. The SARS-CoV-2 pseudovirus was added sequentially to the serum dilutions and incubated at 37°C with 5% CO_2_ supplementation for 60 ± 5 minutes. Serum-virus complexes were then transferred onto plates, previously seeded overnight with Vero E6 cells, and incubated at 37°C and 5% CO_2_ for 20 ± 2 hours. Following this incubation, the luciferase substrate was added to the cells in order to assess the level of luminescence per well. The plates were then read on a luminescence plate reader. The intensity of the luminescence was quantified in relative luminescence units (RLU). The neutralizing titer of a serum sample was calculated as the reciprocal serum dilution corresponding to the 50% neutralization antibody titer (IC_50_) for that sample. The LLOD is 10, which is the first sample dilution (1:10) used as an informative LLOD. LLOQ is based on qualification performed for human samples has been set on 33 IC_50_.

## Wildtype virus neutralization assay (wtVNA)

Leiden University Medical Center (LUMC)assay – Neutralization assays against live SARS-CoV-2 were performed using the microneutralization assay as previously described (Bos et al., 2020), with the modification of a different strain used, SARS-CoV-2 isolate Leiden-0008. Isolate Leiden-0008 (GenBank accession number: MT705206.1) was propagated and titrated in Vero E6 cells using the TCID_50_ endpoint dilution method and the TCID_50_ was calculated by the Spearman-Kärber algorithm as previously described (Hierholzer and Killington, 1996) All work with live SARS-CoV-2 was performed in a BSL3 facility at Leiden University Medical Center. Vero-E6 cells were seeded at 12,000 cells/well in 96-well tissue culture plates one day prior to infection. Heat-inactivated (30 minutes at 56°C) serum samples were analyzed in duplicate. The panel of sera were two-fold serially diluted in duplicate, with an initial dilution of 1:10 and a final dilution of 1:1280 in 60 μL eagle’s minimum essential medium (EMEM) supplemented with penicillin, streptomycin, 2 mM L-glutamine and 2% fetal calf serum (FCS). Diluted sera were mixed with equal volumes of 120 TCID_50_/60 μL Leiden −0008 virus and incubated for 1 h at 37 °C. The virus-serum mixtures were then added onto Vero E6 cell monolayers and incubated at 37 °C in a humidified atmosphere with 5 % CO_2_. Cells either unexposed to the virus or mixed with 120 TCID_50_/60 μL SARS-CoV-2 were used as negative (uninfected) and positive (infected) controls, respectively. At three days post-infection, cells were fixed and inactivated with 40 μL 37% formaldehyde/PBS solution/well overnight at 4 °C. The fixative was removed from cells and the clusters were stained with 50 μL/well crystal violet solution, incubated for 10 minutes and rinsed with water. Dried plates were evaluated for viral cytopathic effect. Neutralization titer was calculated by dividing the number of positive wells with complete inhibition of the virus-induced cytopathogenic effect, by the number of replicates, and adding 2.5 to stabilize the calculated ratio. The neutralizing antibody titer was defined as the log2 reciprocal of this value. A SARS-CoV-2 back-titration was included with each assay run to confirm that the dose of the used inoculum was within the acceptable range of 30 to 300 TCID_50_Public Health England (PHE) assay - Neutralizing antibodies capable of inhibiting wild type virus infections were quantified using the wildtype virus microneutralization assay (MNA) that was developed and qualified for human samples by PHE. The virus stocks used were derived from the Victoria/1/2020 strain (GenBank accession number: MT007544.1).

In brief, 6 two-fold serial dilutions of the heat-inactivated NHP serum samples were prepared in 96-well transfer plate(s). The SARS-CoV-2 wild-type virus was subsequently added to the serum dilutions at a target working concentration (approximately 100 plaque-forming units [Plaque-Forming Unit (PFU)]/well) and incubated at 37°C for 60 to 90 minutes. The serum-virus mixture was then transferred onto assay plates, previously seeded overnight with Vero E6 African green monkey kidney cells and incubated at 37°C and 5% CO_2_ for 60 to 90 minutes before the addition of carboxymethyl cellulose (CMC) overlay medium and further incubation for 24 hours. Following this incubation, the cells were fixed and stained using an antibody pair specific for the SARS-CoV-2 RBD S protein and immunoplaques were visualized using TrueBlueTM substrate. Immunoplaques were counted using an Immunospot Analyzer (Cellular Technology Limited,CTL). The immunoplaque counts were exported to SoftMax Pro (Molecular Devices) and the neutralizing titer of a serum sample was calculated as the reciprocal serum dilution corresponding to the 50% neutralization antibody titer (IC_50_) for that sample.

## Enzyme-Linked Immunospot assay (ELISpot)

IFN-γ /IL-4 Double-Color was performed on freshly isolated PBMCs. PBMC were isolated from ethylene diamine tetraaceticacid (EDTA) whole blood using Ficoll gradient centrifugation (10ml 92% Ficoll-Paque (GE Healthcare) Plus in 1:4 Dulbecco’s phosphate-buffered saline (DPBS)-diluted blood) The ELISpot was performed using the ImmunoSpot Human IFN-γ/IL-4 Double-Color Enzymatic ELISpot Assay Kit according to the manufacturer’s protocol (Cellular Technology Limited). Ethanol-activated 96-well ELISpot plates were coated overnight with anti-human IFN-γ and IL-4 capture antibodies. Cells were plated at a concentration of 250,000 cells per well and stimulated with either cell culture medium in presence of dimethylsulfoxide (DMSO), 2 pools of consecutive 15-mer peptides with 11 amino acid overlap (JPT) spanning the entire length of the SARS-CoV-2 spike protein at a peptide concentration of 2 μg/mL, or 1 μg/mL phytohemagglutinin (PHA) as positive control for 22 hours. Analysis was performed using the CTL ImmunoSpot Analyzer and ImmunoSpot Software (Cellular Technology). Spot-forming units per 1.0 × 10^6^ PBMCs were calculated by subtraction of medium stimulus counts of the individual peptide pools per animal and summed across the 2 peptide pools.

## Intracellular cytokine staining (ICS)

For analysis of intracellular cytokine expression, 1×10^6^ freshly isolated PBMC were stimulated at 37 °C overnight (approximately 15 hours) with either cell culture medium in presence of DMSO, 2 μg/mL SARS-CoV-2 spike protein peptide pools (as described for ELISpot), or 5 μg/mL PHA in the presence of GolgiStop (BD Biosciences). Stimulated cells were first incubated with LIVE/DEAD Aqua viability dye (Thermo Fisher Scientific), followed by surface staining with anti-human monoclonal antibodies CD3-PerCP-Cy5.5 (clone SP34-2, cat. nr. 552852), CD4-APC H7 (clone L200, cat. .nr. 560837), CD8-BV650 (clone SK1, cat. nr. 565289), CD14-BV605 (clone M5E2, cat. nr. 564054), CD69-BV786 (clone FN50, cat. nr. 563834), all from BD Biosciences, and CD20-BV605 (Biolegend, clone 2H7, cat. nr. 302334). Cells were subsequently fixed with Cytofix/Cytoperm buffer (BD Biosciences) and stained intracellularly with anti-human IL-2-PE (clone MQ1-17H12, cat. nr. 560709), IFN-γ-APC (clone B27, cat. nr. 554702) from BD Biosciences, IL-5-Vio515 (clone JES1-39D10, cat. nr. 130-108-099, Miltenyi Biotec), IL-4-PE Dazzle594 (clone MP4-25D2, cat. nr. 500832) and IL-13-BV421 (clone JES10-5A2, cat. nr. 501916), both from Biolegend. Sample acquisition was performed on a LSR Fortessa (BD Biosciences) and data were analysed in FlowJo V10 (TreeStar). Antigen-specific T cells were identified by consecutive gating on single cells (FSC-H versus FSC-A), live cells, size (lymphocytes) (FSC-A versus SSC-A), CD3+, CD4+ or CD8+ cells and CD69+ plus cytokine-positive (the gating strategy is shown in Supplementary Fig. 4). Cytokine-positive responses are presented after subtraction of the background response detected in the corresponding medium stimulated sample of each individual animal. Responders were defined by a technical threshold (Bowyer et al., 2018), the theoretical ability to detect at least 1 event in a cytokine gate and here defined as the reciprocal of the average number of CD4 or CD8 T cells of the medium and peptide pool stimulated samples for each assay run. CD4 Th1 and Th2 T cell subsets were defined by Boolean gating. Th1 subset consists of CD4^+^CD69^+^ T cells expressing IFN-γ and/or IL-2 but not IL-4, IL-5 and IL-13, the Th2 subset was defined as CD4^+^CD69^+^ T cells expressing IL-4 and/or IL-5 and/or IL-13.

## RNA isolation and SARS-CoV-2 subgenomic mRNA assay

RNA was extracted from homogenized lung tissue and from BAL fluid, trachea, and nasal swabs, by use of the QIAamp Viral RNA Mini Kit (Qiagen, Hilden, Germany), according to manufacturer’s instructions. Viral E gene-derived subgenomic messenger RNA (sgmRNA) was quantified using the SuperScript III One-Step RT-PCR System with Platinum Taq DNA Polymerase (Invitrogen, Darmstadt, Germany), with 400 nM concentration of the forward or reverse primer and 200 nM of probe in a 25 μl reaction. The sequence of the sg-leader-specific forward primer, as well as the E-gene specific reverse primer and probe were previously published (Wölfel et al., 2020) Reverse transcription was performed at 50°C for 15 minutes, followed by enzyme activation at 95°C for 2 minutes and 40 PCR cycles of 95°C for 15 seconds and 60°C for 30 seconds. An RNA standard was prepared from a pcDNA3.1 plasmid containing the complete E gene behind the SARS-CoV-2 subgenomic leader sequence (nucleotides. 1-77 of the SARS-CoV-2 genome). Serial dilution of sgm RNA standard with known number of copies was taken along to calculate sgmRNA in copies per ml for swabs or copies per gram tissue. The LLOQs were 2.1×10^3^ copies per mL and 1×10^4^ copies per gram tissue, except for the naïve animals that contributed to the pooled control group, for which the LLOQs were 1.1×10^4^ copies per ml and 5×10^4^ copies per gram tissue.

## Body temperature analysis

A baseline 24-hour body temperature cycle was reconstructed per animal using a multi-day window prior to virus inoculation in which no biotechnical interventions occurred. Fever duration, defined as the net increase in body temperature was calculated as difference relative to the mean of the baseline cycle at corresponding clock times during the first 6 days of the follow-up period. The lower limit of the temperature difference with the baseline was set at zero to reduce the impact of lower body temperatures during daily post-challenge anesthesia and the AUC of the net temperature increase in this period was calculated.

## Lung gross pathology, histopathology and immunohistochemistry

At the end of the follow-up period all animals were necropsied by opening the thoracic and abdominal cavities and all major organs were examined. The extent of pulmonary consolidation was assessed based on visual estimation of the percentage of affected lung tissue. Nasal mucosa, pharynx, trachea, bronchi and all lung lobes were collected for histopathological examination and analysis by immunohistochemistry (IHC). All tissues were immersed in 10% neutral-buffered formalin for fixation, paraffin embedded and stained with haematoxylin and eosin (H&E) for histopathological evaluation. The H&E stained tissue sections were examined by light microscopy (Zeiss Axioplan). For IHC, paraffin sections of all lung lobes were automatically stained (Ventana Discovery Ultra, Roche, France), using rabbit polyclonal anti-SARS-CoV Nucleocapsid protein antibody (NP, Novus NB100-56576). The immunohistochemically stained tissue sections were examined by light microscopy, using a Leica DM2500 light microscope with magnification steps of 25x, 100x, 200x, and 400x.

## Statistical analysis

### ELISA and psVNA

For binding and psVNA neutralizing antibody data, comparisons between specific vaccine groups were made with the two-sample t-test in an analysis-of-variance (ANOVA). Successive time points were compared using the paired t-test per vaccine group. P values were calculated on log10 transformed values.

### wtVNA, ELISpot and ICS

Vaccine groups were compared to the negative control group with the Mann-Whitney U-test. Pairwise comparison between vaccine groups was performed using Tobit ANOVA with vaccine as factor if less than 50% of the titers were at LLOD. The pairwise comparisons between vaccines were done with the z-test. If for an assay any vaccine group had 50% censoring or more, then the pairwise comparisons were done with the Mann-Whitney U-test.

The difference in titer between consecutive time points was calculated per animal for each assay. Depending on the number of censored measurements, the differences were compared with a Tobit ANOVA followed by a post-hoc z test or a sign test.

For all statistical tests the significance level was 5%. No multiple comparison adjustment was applied. All statistical calculations are done in SAS 9.4. (SAS Institute Inc Cary, NC, US).

### Correlation analysis

Correlation analysis between binding antibody concentrations and neutralizing antibody titers or different neutralization assays was calculated using two-sided Spearman rank-correlation.

### Challenge data

Mean nasal and trachea swab area under the curve values of each group were pairwise compared using Tobit ANOVA with post hoc z-test. Mean net temperature difference AUCs were pairwise compared between groups by t-test.

## Notes

### Competing Interest Statement

The authors have declared no competing interest.

### Summary of Updates

Protective efficacy of one- and two-dose Ad26.COV2.S vaccine regimes in aged rhesus macaques Figure 4 and figure 5 are new figures relative to the added efficacy data Supplementary figure 5 and supplementary figure 6 are new figures relative to the added efficacy data

