## Supplementary material for "Immunogenicity and protective efficacy of one- and two-dose regimens of the Ad26.COV2.S COVID-19 vaccine candidate in adult and aged rhesus macaques": Spplemtary Fifure 1 to 6

#### Supplementary figures

##### Supplementary figure 1

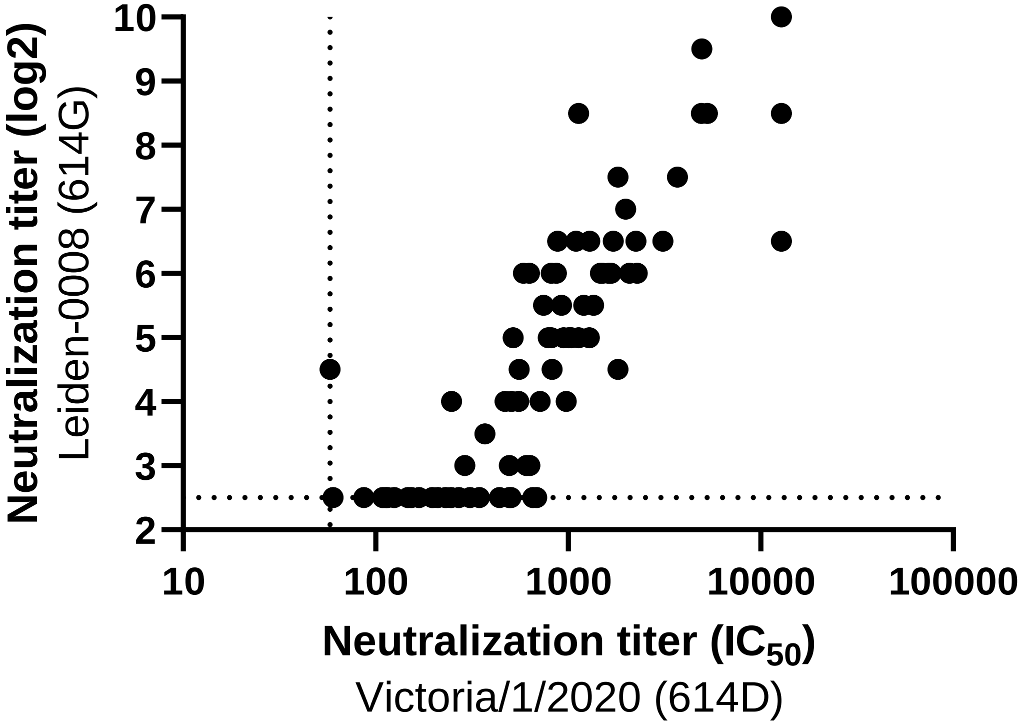

#### Supplementary figure 2

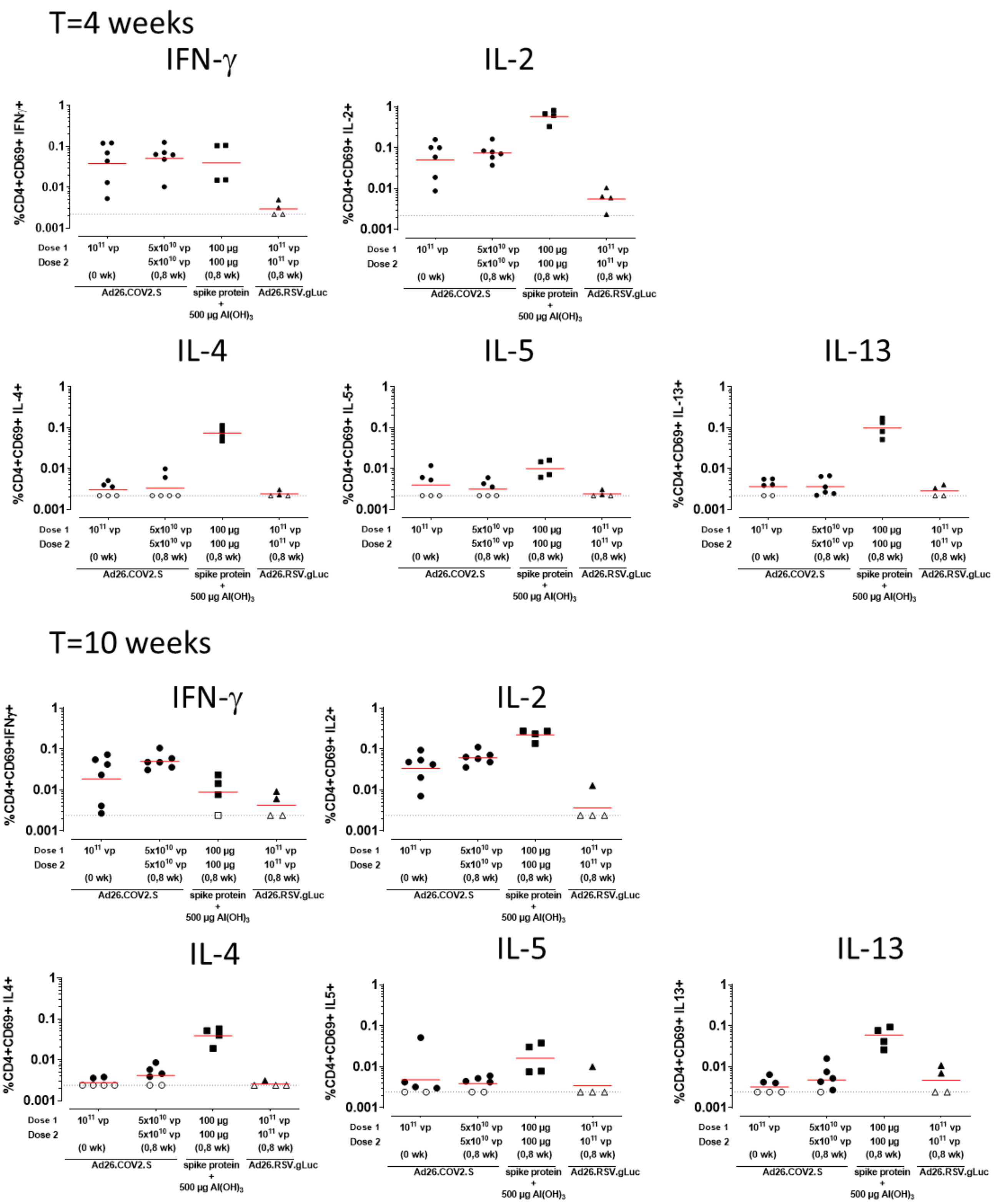

#### Supplementary figure 3

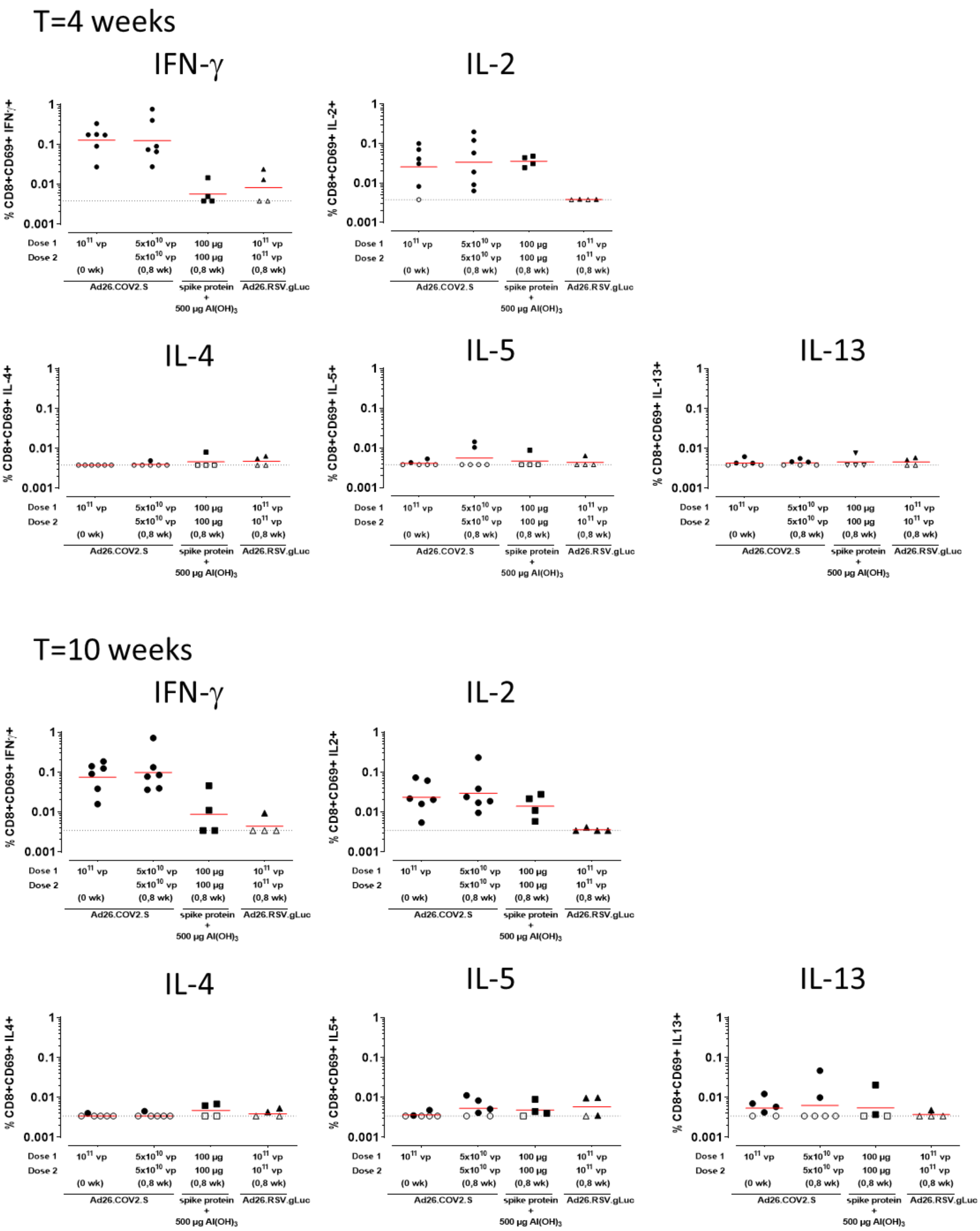

### Supplementary figure 4

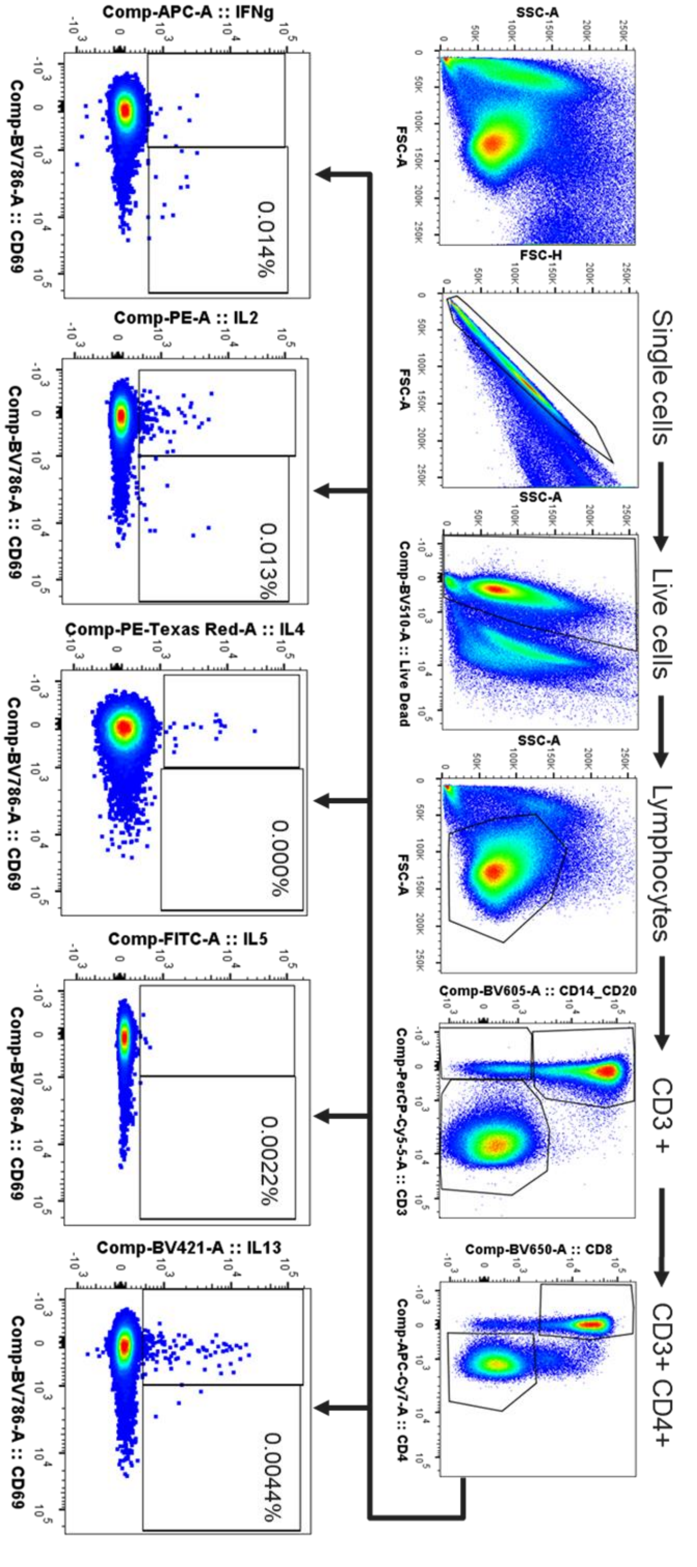

Supplementary figure 5

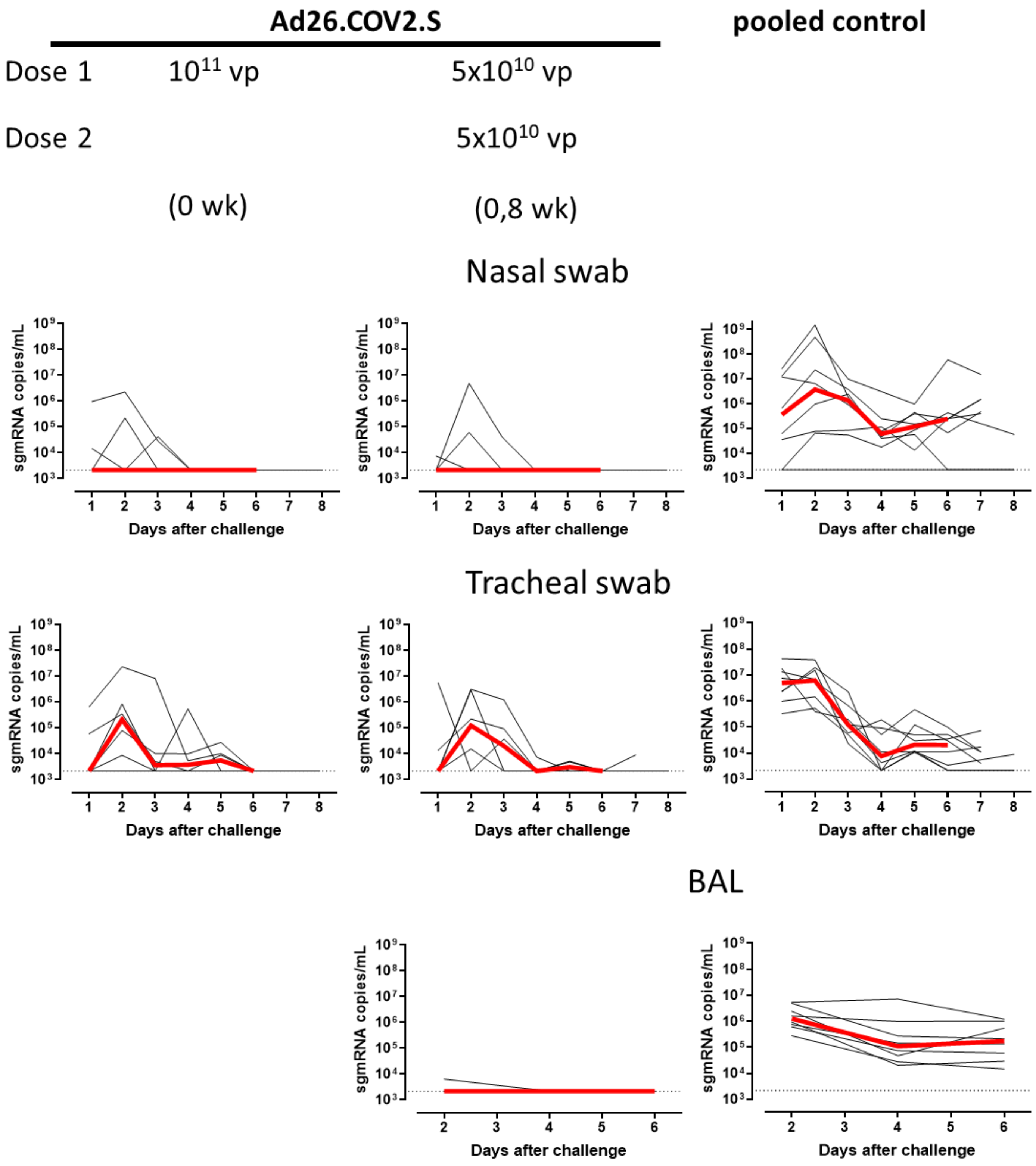

#### Supplementary figure 6

**A**

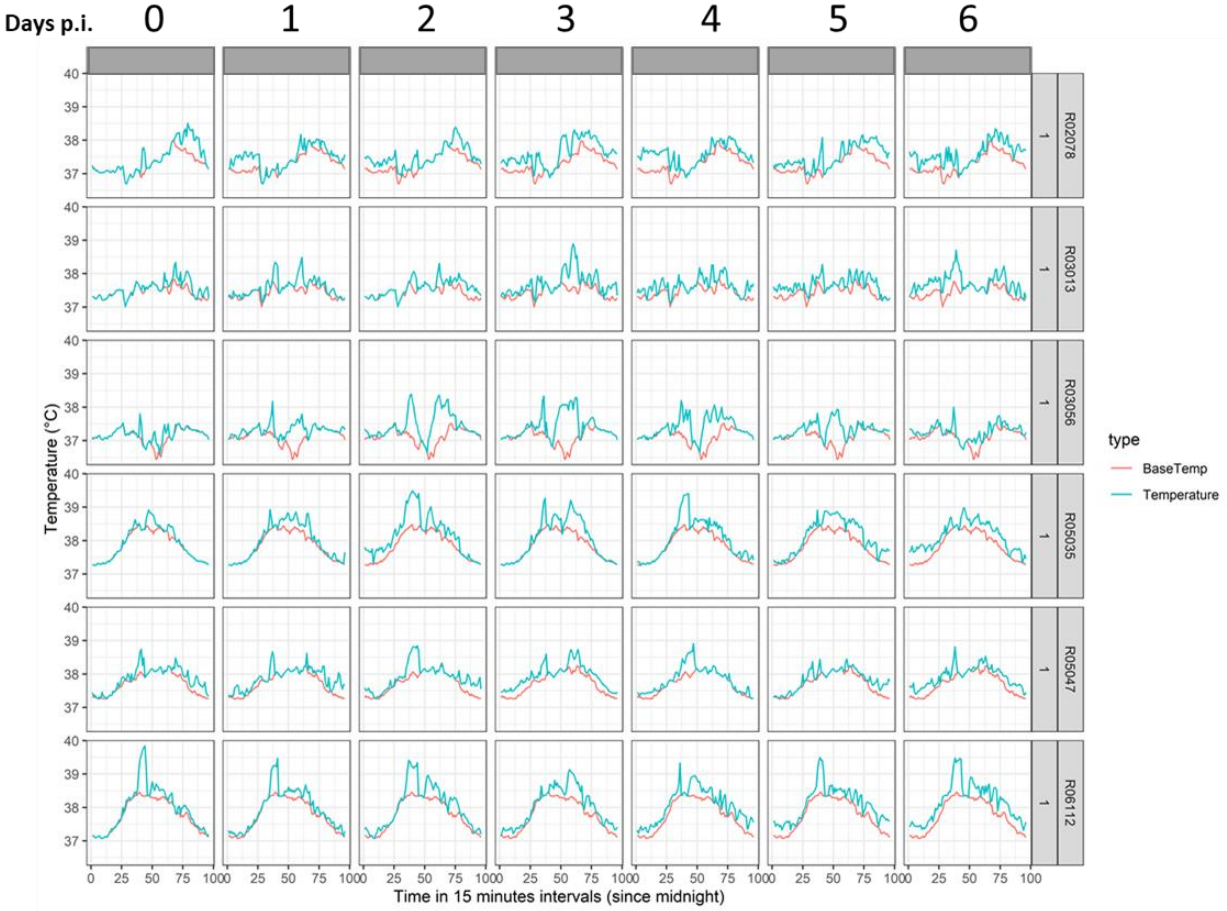

**B**

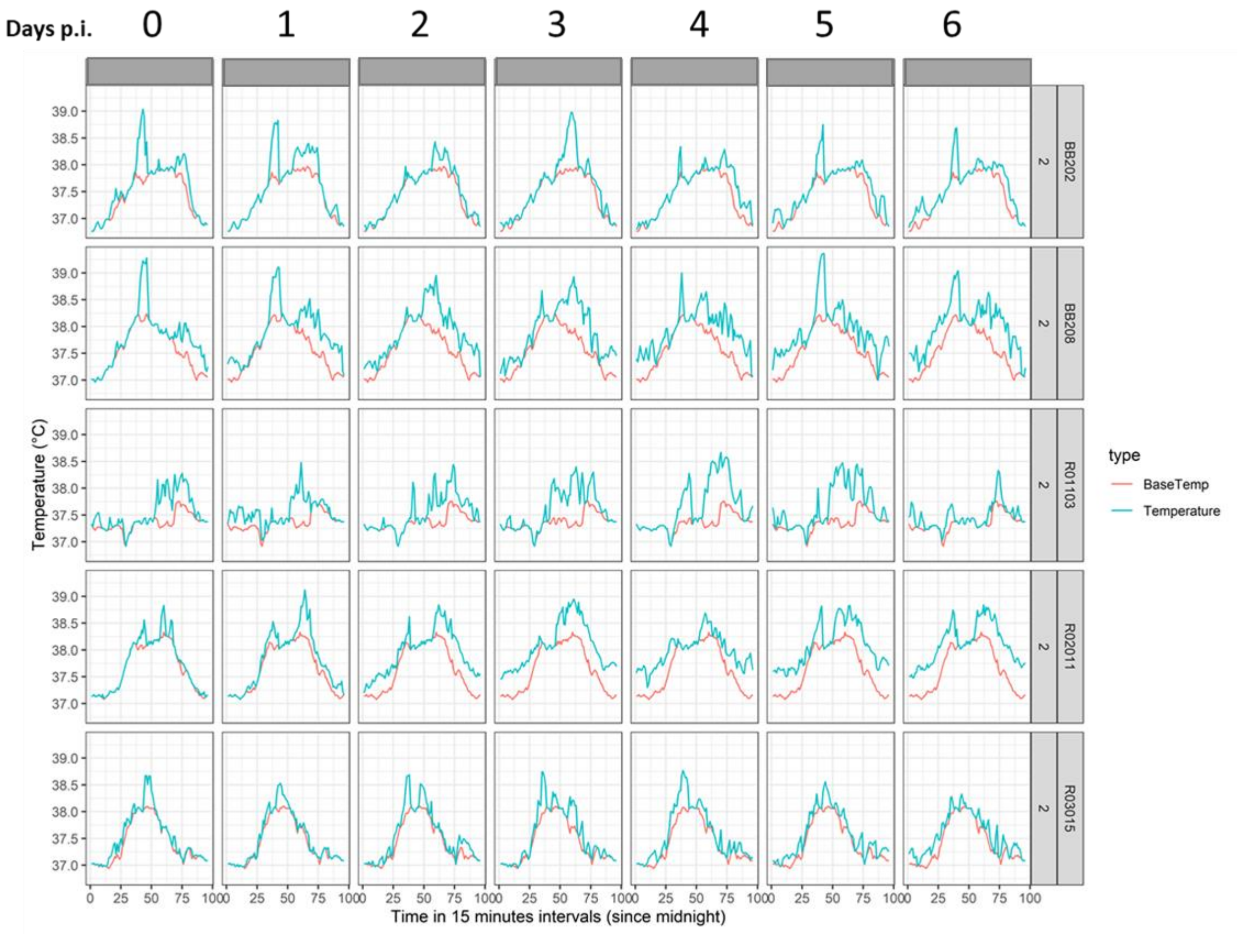

Days p.i.
0
1
2
3
4
5
6
BB202
BB208
R01103
R02011
R03015
2
39.0
38.5
38.0
37.5
37.0
Temperature (°C)
0
25
50
75
100
Time in 15 minutes intervals (since midnight)
type
BaseTemp
Temperature

**C**

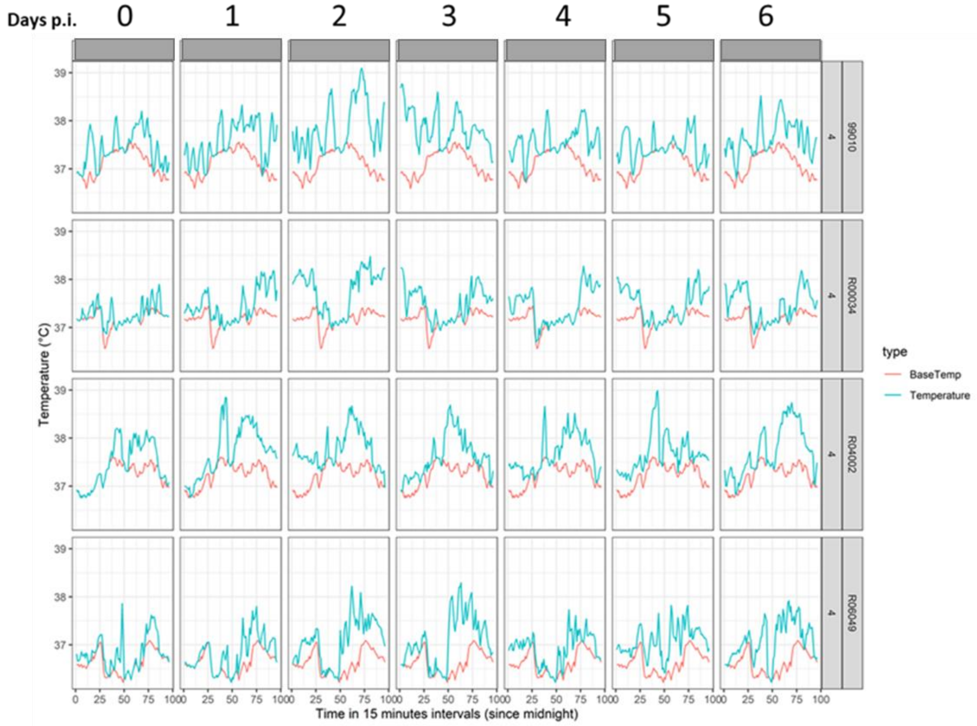

Days p.i.
0
1
2
3
4
5
6
99010
R00034
R04002
R06049
4
Temperature (°C)
Time in 15 minutes intervals (since midnight)
type
BaseTemp
Temperature


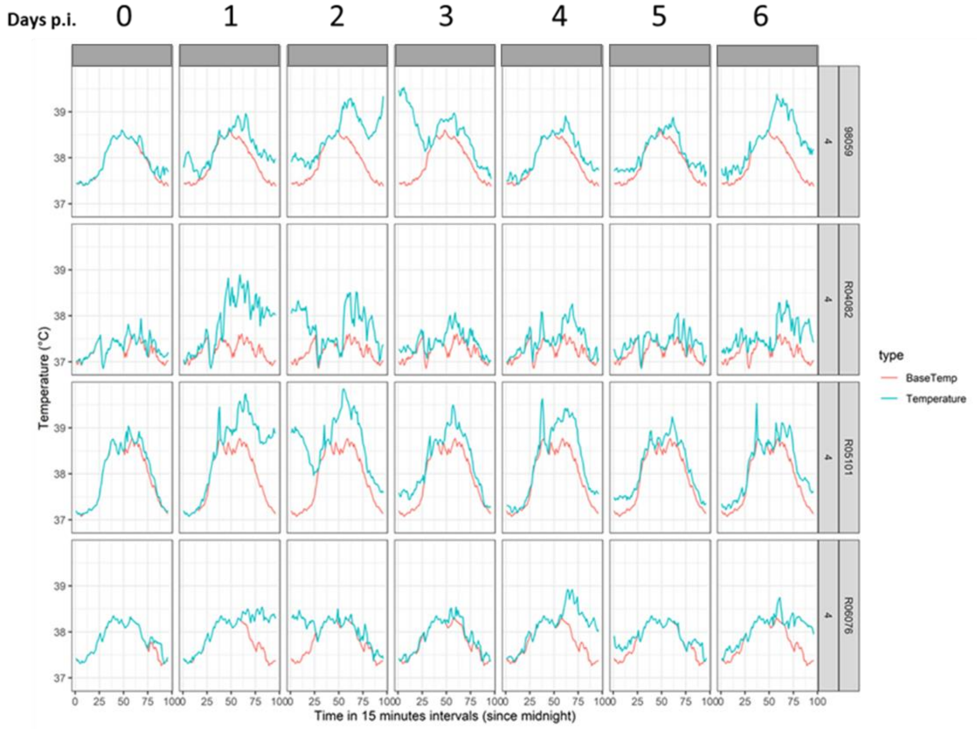

Days p.i.
0
1
2
3
4
5
6
98059
R04082
R05101
R06076
4
Temperature (°C)
Time in 15 minutes intervals (since midnight)
type
BaseTemp
Temperature

#### Supplementary figure legends

**Supplementary figure 1**

Correlation between SARS-CoV-2 neutralizing antibody titers using the Leiden-0008 strain (LUMC) and the Victoria/1/202 strain (PHE) per animal for all groups and timepoints except the sham control group and week 0. The dotted lines indicate the LLOD for each assay.

**Supplementary figure 2**

Spike protein-specific T cell responses as measured by intracellular cytokine staining at indicated timepoints. Frequency of $CD4^{+}CD69^{+}$ T cell expressing cytokines. Gating strategy is provided in supplemental figure 4. The geometric mean response per group is indicated with a horizontal line. The dotted line indicates the technical threshold. Open symbols denote samples at technical threshold.

**Supplementary figure 3**

Spike (S) protein-specific T cell responses as measured by intracellular cytokine staining at indicated timepoints. Frequency of $CD8^{+}CD69^{+}$ T cell expressing cytokines. Gating strategy is provided in supplemental figure 4. The geometric mean response per group is indicated with a horizontal line. The dotted line indicates the technical threshold. Open symbols denote samples at technical threshold.

**Supplementary figure 4**

Intracellular cytokine staining gating strategy to identify cytokine-expressing spike protein-specific CD4+-and CD8+ T cells. Frequencies denote frequency of parent gate.

**Supplementary figure 5**

Viral load (sgmRNA copies/mL) kinetics in swabs and BAL after SARS-CoV-2 inoculation of vaccinated aged rhesus macaques. Black lines represent individual animals, red lines the group median up to day 6, the last day of follow-up prior to scheduled euthanasia. Dashed horizontal line the LLOQ. Data of a related challenge of naïve animals (n=4) using an identical challenge strain, challenge regimen and readouts were added to the sham control group data, collectively referred to as pooled control, to increase statistical power. Due to small differences in the LLOQ of the sgmRNA assay between the 2 studies, five trachea swab samples of the pooled control are set to the higher LLOQ.

**Supplementary figure 6**

Body temperatures after SARS-CoV-2 inoculation of vaccinated aged rhesus macaques on day of infection and for 6 consecutive days afterwards. Calendar dates are on top, animal id and group number labels on the righthand side. Orange lines indicate the daily baseline temperature profile derived from a multi-day window prior to virus inoculation in which no biotechnical interventions occurred. The blue line is the temperature of each animal post infection. A) body temperatures of the $1x10^{11}$ vp Ad26.COV2.S group B) body temperatures of the $5x10^{10}$ vp Ad26.COV2.S group. C) body temperatures of the pooled control animal group. p.i.; post-infection.
